# Divide and conquer – machine-learning integrates mammalian, viral, and network traits to predict unknown virus-mammal associations

**DOI:** 10.1101/2020.06.13.150003

**Authors:** Maya Wardeh, Marcus S.C. Blagrove, Kieran J. Sharkey, Matthew Baylis

**Affiliations:** Department of Livestock and One Health, Institute of Infection, Veterinary & Ecological Sciences, University of Liverpool, Liverpool Science Park IC2 Building, 146 Brownlow Hill, Liverpool L3 5RF; Department of Evolution, Ecology and Behaviour, Institute of Infection, Veterinary & Ecological Sciences, University of Liverpool, Merseybio Building, Crown Street L69 7ZB; Department of Mathematical Sciences, University of Liverpool, Peach Street, Liverpool, L69 7ZL

## Abstract

Our knowledge of viral host ranges remains limited. Completing this picture by identifying unknown hosts of known viruses is an important research aim that can help identify zoonotic and animal-disease risks. Furthermore, such understanding can be used to mitigate against viral spill-over from animal reservoirs into human population. To address this knowledge-gap we apply a divide-and-conquer approach which separates viral, mammalian and network features into three unique perspectives, each predicting associations independently to enhance predictive power. Our approach predicts over 20,000 unknown associations between known viruses and mammalian hosts, suggesting that current knowledge underestimates the number of associations in wild and semi-domesticated mammals by a factor of 4.3, and the average mammalian host-range of viruses by a factor of 3.2. In particular, our results highlight a significant knowledge gap in the wild reservoirs of important zoonotic and domesticated mammals’ viruses: specifically, lyssaviruses, bornaviruses and rotaviruses.

## INTRODUCTION

Thousands of viruses are known to affect mammals, with recent estimations indicating that less than 1% of mammalian viral diversity has been discovered to date^1^. Some of these viruses have a very narrow host range, whereas others such as rabies and West Nile viruses have very wide host ranges (rabies can theoretically infect any mammal^2^). Host range is an important predictor of whether a virus is zoonotic^3^ and therefore pose risk to humans. For example, Severe acute respiratory syndrome-related (SARS) and Middle East respiratory syndrome-related (MERS) coronaviruses are both believed to have originated in bats, but through a host range that includes other mammals (e.g. palm civets^4^, camels^5^) they have successfully infected humans. Most recently, the emergence of SARS-CoV2 has been linked to spread from bats to Malayan pangolins^6^. Knowing the potential host range of viruses is essential to our efforts to develop preventive and protective strategies and policies.

However, our knowledge of the host range of viruses remains limited^1,3,7^ and the knowledge we have is hugely biased towards humans and domesticated mammals. For example, there is a significant gap between the number of known human viruses (274 species^8^), and those of wild primates (e.g. only 5 species in the toque macaque - *Macaca sinica*^8^) which is largely a result of differential research effort. Surveillance and research efforts often intensify during and after significant outbreaks, leading to further biases; for instance, recent efforts to identify potential reservoirs of SARS-Cov-2 have led to the identification of two new virus species in wild pangolins (*Manis javanica* and *Manis pentadactyla*)^9^, and a pangolin coronavirus^6^, thereby doubling the number of known viruses of pangolins.

Despite these biases, the knowledge accumulated so far provides a valuable resource which can be exploited to estimate the extent of which we are under-observing associations between known viral agents and mammalian hosts. Here, we integrate this knowledge into a machine-learning driven framework to predict unknown (i.e. either potential or undocumented/unobserved) associations between known viruses and their mammalian hosts. The novelty of our framework lies in its multi-perspective approach whereby each possible virus-mammal association is predicted three times: 1) from the perspective of the mammalian hosts (e.g., based on the traits of the viruses known to infect wildcats - *Felis silvestris*, which other known viruses could also infect them); 2) the perspective of the viruses (e.g. based on the traits of mammalian species in which West Nile virus has been found, which other mammals can carry this virus ?); and 3) from the perspective of the network linking known viruses with their mammalian hosts.

This network presents a global view of how these viruses are shared between mammalian hosts. This sharing exhibits certain characteristics (e.g. DNA vs RNA viruses^10^; bats vs rodents^11^) which could only be captured at the global level. Various network topological features have been exploited to provide significant insight into patterns of pathogen sharing^11^, disease emergence and spill-over events^12^, and as means to predict missing links in a variety of host-pathogen networks^13^ including helminths^14^, and viruses^15^. Here we express the topology of our virus-mammal network in terms of counts of *potential motifs*^16^. Motifs^17^ are miniature subgraphs which constitute the building blocks of larger, more complex networks^18^. Motifs express specific functions or topological features of the underlying network, and have been used to capture complex and indirect interactions in a variety of systems including biology^19–21^, ecology^22,23^ and disease emergence^24^.

Our framework utilises 6,331 associations between 1,896 viruses and 1,436 terrestrial mammals, representing 0.23% of all possible associations between our mammals and their viruses. It assesses how much these associations are underestimated by predicting which unknown species-level associations are likely to exist in nature (or do already exist but yet undocumented). We aggregate these predictions to enhance estimation of the host-range of known mammalian viruses, and to highlight variation in the degree of underestimation at the level of mammalian order (particularly in wild and semi-domesticated species), and viral group (Baltimore classification), family, and genus. In addition, we highlight significant knowledge gaps in mammalian reservoirs of known zoonoses and equivalent viruses in important domesticated mammals. By investigating this underestimation from three separate points of view, we enhance the overall predictive performance and capture local (at the level of a single viral or mammalian species), as well as global (aggregated) variations in our knowledge gaps.

## RESULTS

Our framework to predict unknown associations between known viruses and their mammalian reservoirs comprised three distinct perspectives: viral, mammalian and network. Each perspective produced predictions from a different vantage point (that of each virus, each mammal, and the network connecting them respectively). Subsequently, their results were consolidated via majority voting. This approach suggested that 20,832 (median, 90% CI = [2736, 97062], hereafter values in square brackets represent 90% CI) unknown associations potentially exist between mammals and known viruses, (18,920 [2440, 91517] in wild or semi-domesticated mammals). Our results indicated *a* ∼*4*.*29-fold increase (*[∼1.43, ∼16.33]) in virus-mammal associations (∼4.89 [∼1.5, ∼19.81] in wild and semi-domesticated mammals).

We validated our framework by training its constituent models using a stratified random subset of our input data (2,315,391 known and unknown virus-mammal associations), and then estimated its performance metrics against the remaining data (408,701 instances). Our framework achieved an overall AUC = 0.956, F1-Score = 0.310, and TSS = 0.915 when evaluated against the held-out test data (Supplementary Note 5). Additionally, we conducted a systematic test to predict removed virus-mammal associations. In this test, we systematically removed one known virus-mammal association at a time from our framework, recalculated all models input (including from network) and attempted to predict these removed associations. Our framework succeeded in predicting 90% of removed associations (90.70% for associations removed for viruses, 89.92% for associations removed from mammals, Supplementary Results 1).

In the following subsections we first illustrate the mechanism of our framework via an example, then further explore the predictive power of our approach for viruses and mammals.

### Example

Our multi-perspective framework generates predictions for each known or unknown virus-mammal association (2,722,656 possible associations between 1,896 viruses and 1,436 terrestrial mammals). We highlight this functionality using two examples (Figure 1): West Nile Virus (WNV) a flavivirus with wide host range, and the bat *Rousettus leschenaultia* (order: Chiroptera). We first consider each of our perspectives separately, and then showcase how these perspectives are merged to produce final predictions.

**Figure 1.**
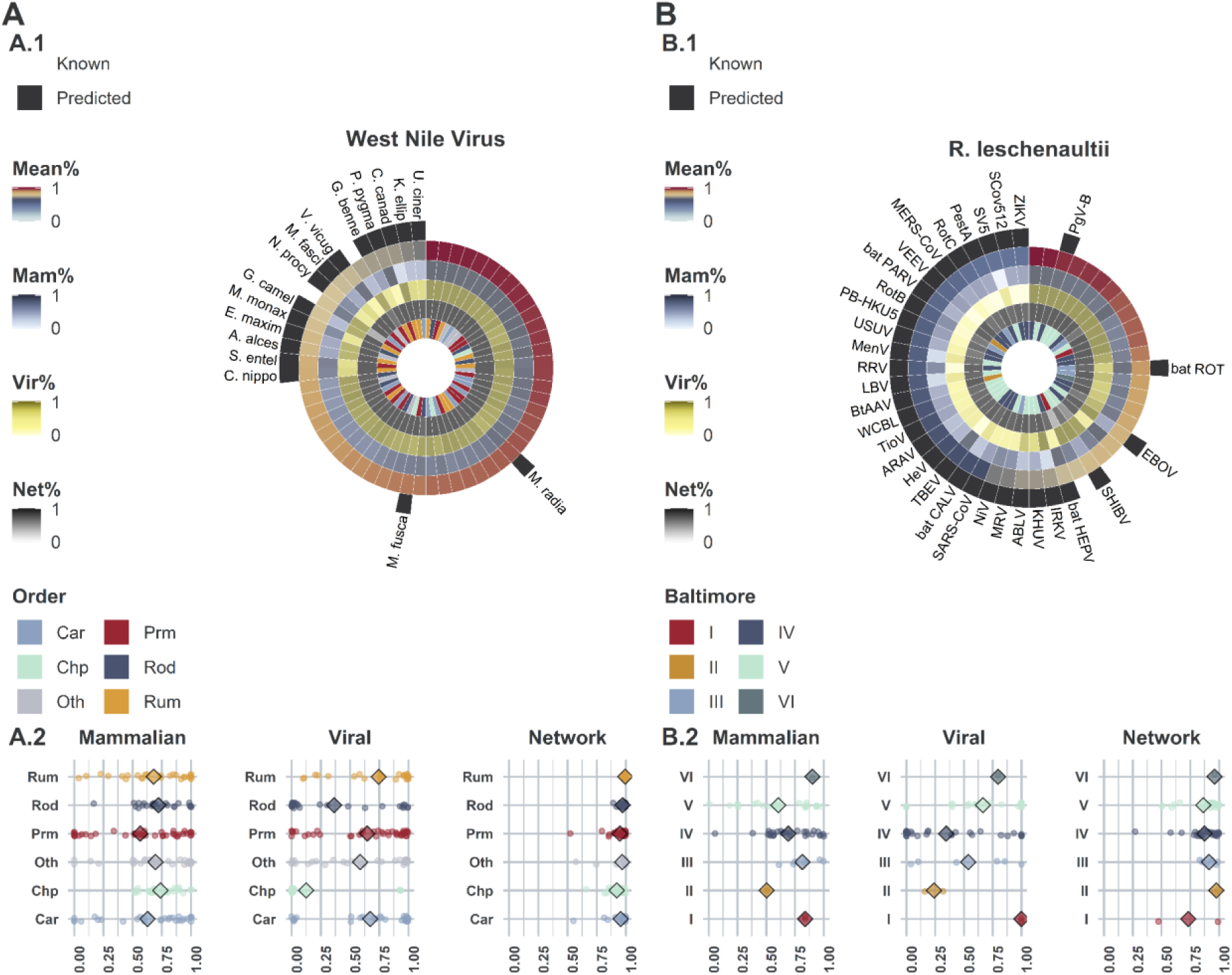
Example showcasing final and intermediate predictions of West Nile Virus (WNV), and *Rousettus leschenaultii*. **Panel A: Results for WNV. Panel A.1: Top 60 predicted hosts of WNV.** Host species were ordered by mean probability of predictions, and top 60 were selected. Circles represent the following information in order: 1) whether association is known (found in EID2) or not (potential or undocumented). Host species are omitted for known associations, full results available in supplementary results. 2) Mean probability across three perspectives. 3) Probabilities of associations between WNV and mammals derived from mammalian perspective. These probabilities are obtained from 60 models, one for each predicted host, trained with viral features, instead of absolute classification (yes or no). 3) Probabilities of associations between WNV and mammals derived from viral perspective (WNV model trained with mammalian features). 4) Probabilities of associations between WNV and mammals as derived from the network perspective (trained with potential-motifs as features). 6) Taxonomic order of predicted hosts. **Panel A**.**2: probability of WNV-mammal associations in the three perspectives per host order**. Supplementary tables S10 presents full results. **Panel B: Results for *R. leschenaultii*. Panel B**.**1: top 50 predicted viruses of R. leschenaultii**. Virus species were ordered by mean probability of predictions derived from mammalian, viral and network perspectives, and top 50 were selected. Circles as per Panel A.1. Baltimore represents Baltimore classification (7 groups: I-VII). **Panel B**.**2: probability of virus-*R. leschenaultii* associations in the three perspectives per Baltimore group**. Supplementary tables S11 presents full results.

#### The mammalian perspective

our mammalian perspective models, trained with features expressing viral traits (table 1), suggested a median of 90 [17, 410] unknown associations between WNV and terrestrial mammals could form when predicting virus-mammal associations based on viral features alone - ∼2.61 fold increase [∼1.3, ∼8.32]. Similarly, our results indicated that 64 [4, 331] new associations could form between our selected host (*R. leschenaultia*) and our viruses (∼4.37-fold increase [∼1.21, ∼18.42] - *Supplementary Results 2*).

**Table 1.** Viral traits & features used to build our mammalian models (N=699). Host-driven features were calculated for each virus-mammal association (N=1,896 x 1,436 = 2,722,656) Full details of these features, their sources and full justification are listed in Supplementary Note 2.

| Category | Viral Feature | Data type | Reason for inclusion |
| --- | --- | --- | --- |
| **Host-driven** | Mean phylogenetic distance between hosts | Continuous | Capturing phylogenetic and ecological distances between each virus' known hosts and each mammal in our study. |
|  | Mean ecological distance between hosts |  |  |
|  | maximum phylogenetic breadth <sup>25</sup> |  | Greater phylogenetic breadth indicates more generalist potential of the virus. |
| **Virus genome & capsid** | RNA | Binary | RNA viruses mutate/adapt faster <sup>26</sup>, and are generally deactivate quickly when exposed to the environment. |
|  | retro-transcribing |  | Retroviruses are generally very conserved <sup>27</sup>, have to enter the nucleus <sup>28</sup> and insert into the genome. Additional steps may require specificity and limit range. |
|  | Negative sense/positive sense |  | Sense affects replication cycle and range of host enzymes needed. |
|  | Circular/linear |  | Circular/linear genome affects enlisting host enzymes for replication and translation <sup>29</sup>. |
|  | Monopartite/segmented |  | Segmented viruses can undergo recombination if two strains of the same virus infect a cell <sup>30</sup>. This can lead to host range changes of segments of the genome. |
|  | Enveloped |  | Envelopes are derived from the host cell membrane, so can affect specific-host immune activation. Enveloped viruses deactivate rapidly in the external environment (often requiring direct transfer). The envelope will change upon infection of a new host<sup>31</sup>. |
|  | GC-content | Continuous | High GC content usually leads to higher thermostability of the genome <sup>32</sup>. |
|  | Genome size |  | Genome size is indicative of many aspects of the virus such as complexity, DNA/RNA, and replication type. |
| **Virus replication, release and cell entry** | Cytoplasm | Binary | Replication site is linked to RNA/DNA genome – if a virus has a DNA stage it must replicate in the nucleus and overcome additional cell barriers. |
|  | Release | Categorical | Affects rate of virus production, cell life-span and means of presentation to the immune system<sup>33</sup>. |
|  | Cell entry |  | Availability of receptors influences potential host range. |
| **Transmission routes** | 8 main transmission routes | Binary for each route | Route(s) of transmission affected by structure/stability of virus and nature of interaction between potential hosts. |

#### The viral perspective

our viral models, trained with features expressing mammalian traits, phylogeny, and spatial distribution (table 2), indicated a median of 48 [0, 214] new hosts of WNV (∼1.86-fold increase [∼1, 4.82]). Results for our example host (*R. leschenaultia*) suggested 18 [3, 76] existing viruses could be found in this host (∼1.95-fold increase [∼1.16, ∼5.00]) - *Supplementary Results 3*).

**Table 2.** mammalian traits & features used to build our viral models (N=556). Host-driven features were calculated for each virus-mammal association (n=1,896 x 1,436 = 2,722,656) and were. Full details of these features, their sources and full justification are listed in Supplementary Note 3.

| Category | Mammalian Feature | Reason for inclusion |
| --- | --- | --- |
| **Phylogeny** | Mean phylogenetic distance to known hosts. | Linked to sharing of viruses between mammals<sup>25,34,35</sup>. |
|  | Evolutionary distinctiveness | Can correlate negatively with pathogen species richness<sup>36</sup>. |
| **Taxonomy & domestication** | Order & family | Can affect host-pathogen<sup>37</sup>, particularly viral, associations<sup>25</sup>. |
|  | Domestication | Might influence sharing of viruses between host groups. Domesticated mammals and human might share more viruses with each other than related wild species. |
| **Ecological traits** | Morphological traits (body mass) | A key feature in terms of metabolism and adaption to environment. |
|  | Life-history traits (Maximum age, age at sexual maturity, activity cycle, and migration) | Potentially relevant in terms of within-host dynamics of viruses. |
|  | Reproductive traits (gestation period length, litters per year, litter size and weaning age) |  |
|  | Habitat utilisation | Similar habitat utilisation might correlate with contact with similar viruses. |
|  | Diet (proportional use of 10 categories) | Similar dietary habit might associate with similar viral assemblage. |
|  | Mean ecological distance | Indicates if a potential host species is *ecologically* close to or distant from the virus' preferred host range. We based this distance on a generalised form of Gower's distance matrices<sup>38,39</sup> incorporating all ecological traits. |
| **Geo-spatial features** | Area size | Might lead to exposure to larger number or more diverse viruses. |
|  | Climate (mean temperature & precipitation) | Climate has been shown to influence a number of human and domestic mammal pathogens (including viruses)<sup>40,41</sup>. |
|  | Natural land cover diversity / Agriculture and farming diversity | These factors have been found to influence certain categories of host-pathogen associations<sup>42</sup>. Supplementary Note 3 lists further details of mammalian geo-spatial feature extraction. |
|  | Mammalian biodiversity |  |
|  | Urbanisation / human population |  |

#### The network perspective

our network models indicated a median of 721 [448, 1317] (∼13.88 [9, 24.52] fold increase) unknown associations between WNV and terrestrial mammals, and that 246 [91, 336] existing viruses could be found in our selected host (*R. leschenaultia*), equivalent to ∼13.95 [∼5.79, ∼18.68] fold increase (*Supplementary Results 4*).

#### Consolidation of perspectives

Considering that each of the above perspectives approached the problem of predicting virus-mammal associations from a different angle, the agreement between these perspectives varied. In the case of WNV: mammalian and viral perspectives achieved 92.3% agreement [72.6% – 98.5%] (Cohen’s kappa, applied to both positive and negative links); mammals and network perspectives had 55.3% agreement [33.4% - 69.5%]; and viruses and network had 52.9% agreement [19.8% – 68.7%]. In the case of *R. leschenaultia* these numbers were as follows: 96.15% [82.44%, 99.58%], 87.24% [76.37%, 95.04%], and 87.61% [75.90%, 95.25%], respectively. The agreements between our perspectives across the 2,722,656 possible associations were as follows: 98.04% [90.36%, 99.73%] between mammalian and viral perspectives, 96.71% [88.62%, 98.92%] between mammalian and network perspectives, and 97.11% [91.57%, 98.95%] between viral and network perspectives.

#### After voting

our framework suggested that a median of 117 [15, 509] new or undetected associations could be missing between WNV and terrestrial mammals (∼3.45-fold increase [∼1.3, ∼12.2]). Similarly, our results indicated that *R. leschenaultia* could be susceptible to an additional 45 [5, 235] viruses that were not captured in our input (∼1.37-fold increase [∼1.26, ∼13.37]). Figure 1 illustrates top predicted and detected associations for WNV and *R. leschenaultia*.

### Mammalian host range of viruses

#### Relative importance of viral features

our multi-perspective approach enabled us to assess the relative importance of viral traits (Table 1) to each of our mammalian models. This in turn showcased variations of how these viral traits contribute to the models at the level of individual species (e.g. humans), and at an aggregated level (e.g. by order or domestication status). The results are highlighted in Figure 3 (panel A). Our results indicate that mean phylogenetic (median = 95.4% [75.6%, 100%]) and mean ecological (90.90% [43.50%, 100%]) distances between potential and known hosts of each virus are the top predictors of associations between the focal host and each of the input viruses. Maximum phylogenetic breadth was also important (74.7 0%, [16.60%, 100%]).

**Figure 2.**
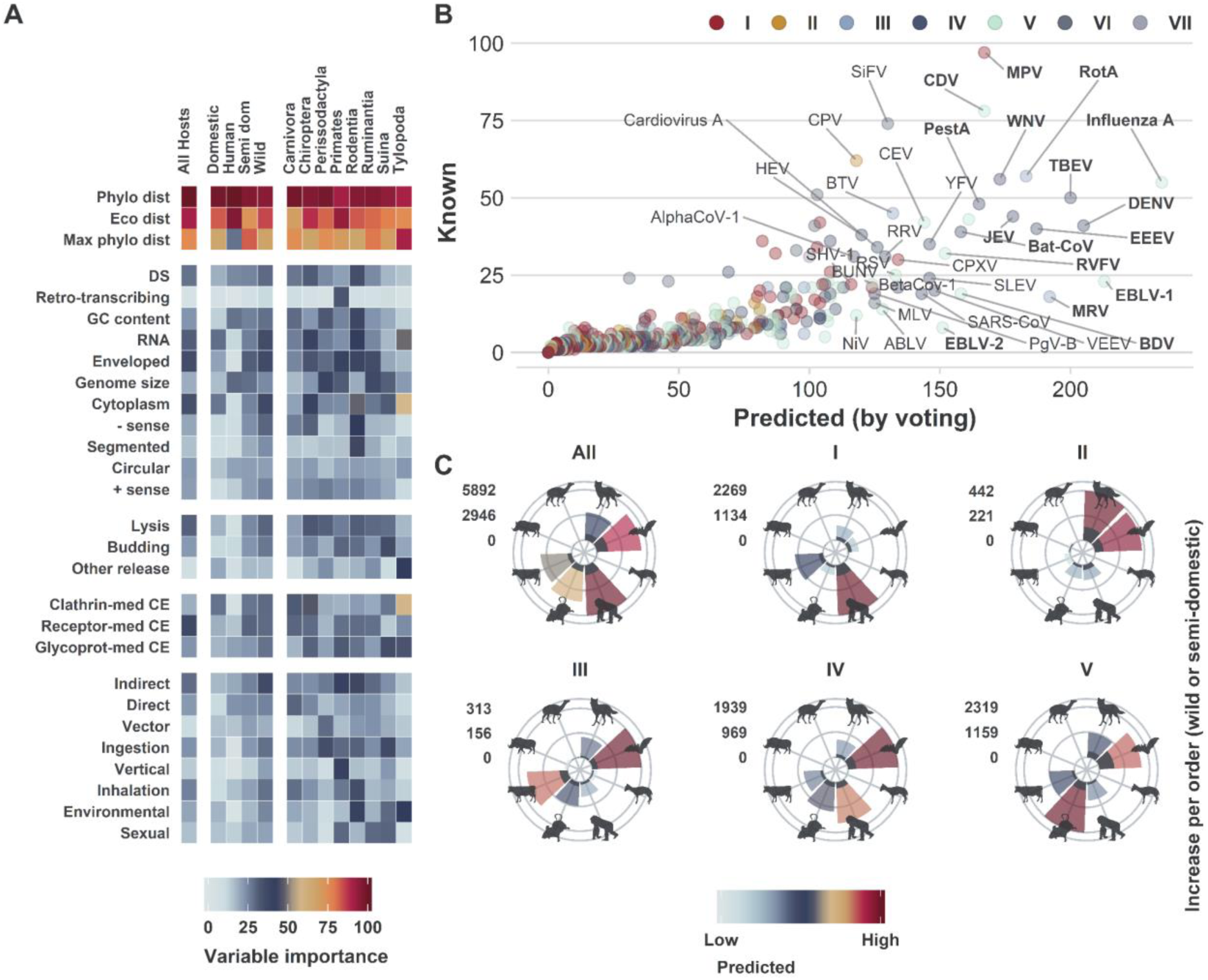
Results (viruses). **Panel A – Variable importance (relative contribution) of viral traits to mammalian perspective models.** Variable importance is calculated for each constituent model (N=699) of our mammalian perspective (trained with viral features), and then aggregated (median) per each reported group (columns). **Panel B – changes in mammalian host range of viruses from known (recorded in EID2) to predicted (by voting)**. Rabies lyssavirus was excluded from panel B to allow for better visualisation. Top 40 (by number of predicted mammalian hosts) are labelled. Species in bold have over 150 predicted hosts (Supplementary table S12 lists details of these viruses including CI). **Panel C – predicted virus-mammal associations (per mammalian order) between viruses and wild and semi-domesticated mammals**. Following orders (clockwise) are presented: Carnivora, Chiroptera, Perissodactyla, Primates, Rodentia, Ruminantia, Suina and Tylopoda. Dark grey bars indicate detected associations. Source of the silhouette graphics is PhyloPic.org. (Supplementary table S13 lists results per mammalian order).

**Figure 3.**
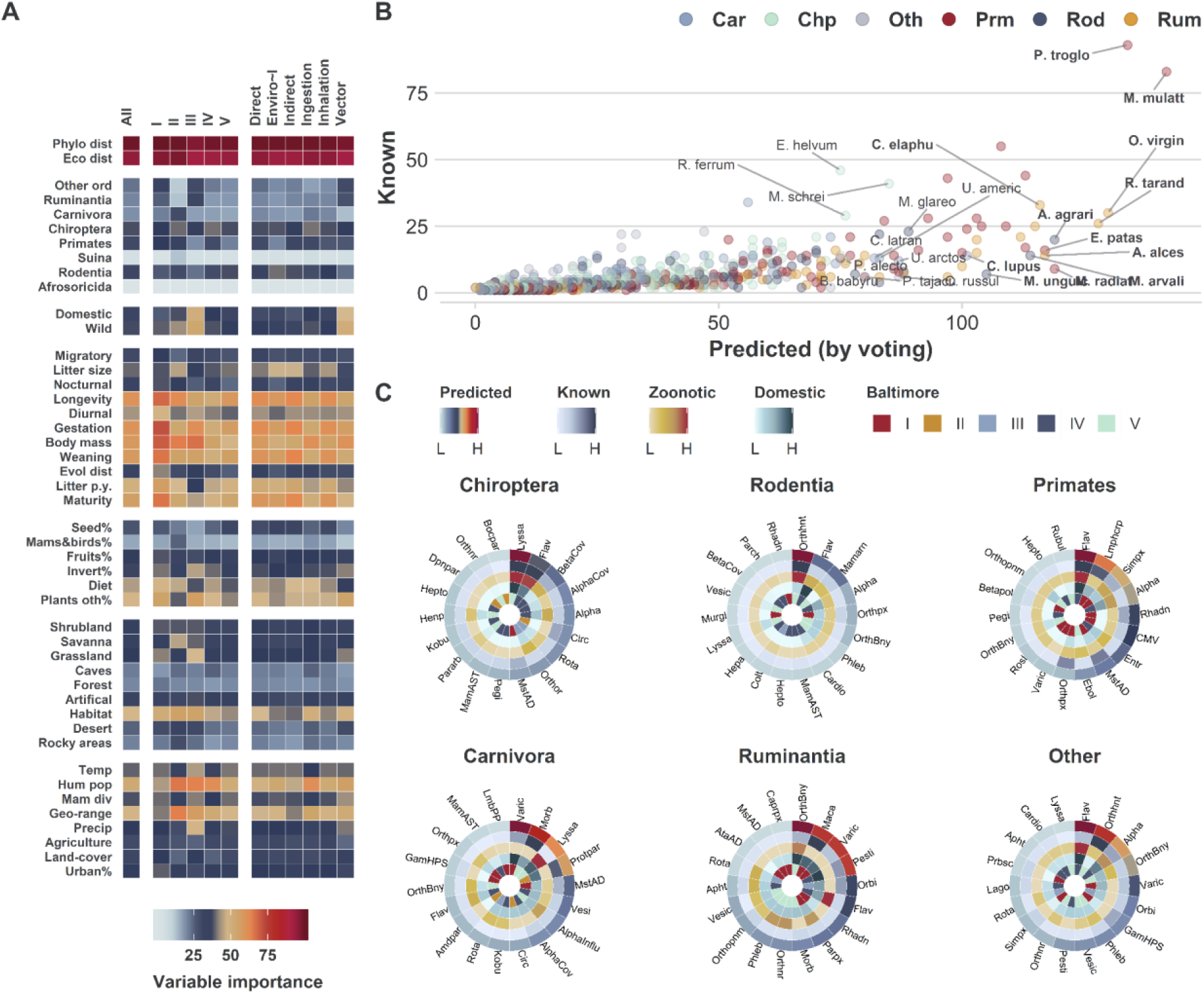
Results (Mammals). **Panel A – variable importance (relative contribution) of mammalian traits to viral perspective models.** Variable importance is calculated for each constituent model (N=556) of our viral perspective (trained with mammalian features), and then aggregated (median) per each reported group (columns). **Panel B – changes in number of viruses per wild or semi-domesticated mammalian host species from known (recorded in EID2) to predicted (by voting)**. Labelled mammals are as follows: top 4 (by number of predicted viruses) for each of carnivora, Chiroptera, Primates, Rodentia, and Ruminantia; and top 3 of other orders. Species in bold have 100 or more predicted viruses (Supplementary table S14). **Panel C – Top 18 genera (by number of predicted wild or semi-domesticated mammalian host species) in selected orders (**Other indicated results for all orders not included in the first five circles). Each order figure comprises the following circles (from outside to inside): 1) Number of hosts predicted to have an association with viruses within the viral genus. 2) Number of hosts detected to have association. 3) Number of hosts predicted to harbour viral zoonoses (i.e. known or predicted to share at least one virus species with humans). 4) Number of hosts predicted to share viruses with domesticated mammals of economic significance (domesticated mammals in orders: Carnivora, Lagomorpha, Perissodactyla, Ruminantia, Suina and Tylopoda). 5) Baltimore classification of the selected genera. (Supplementary table S15 lists full results per mammalian order and virus genus).

#### Mammalian host range

Our results suggested that the average mammalian host range of our viruses is 14.33 [4.78, 54.53] (average fold increase of ∼3.18 [∼1.23, ∼9.86] in number of hosts detected per virus). Overall, RNA viruses had the average host range of 21.65 [7.01, 82.96] hosts (∼4.00-fold increase [∼1.34, ∼14.15]). DNA viruses, on the other hand, had 7.85 [2.81, 29.47] hosts on average (∼2.43 [∼1.14, ∼6.89] fold increase). Table 3 lists the results of our framework at Baltimore group level and selected family and transmission routes of our viruses. Figure 2 illustrates predicted mammalian host range of our viruses (panel B), and the increase in host range of viruses in species-rich mammalian orders of interest (panel C).

**Table 3.** Predicted host-range of viruses per Baltimore group, family (top 15 families, ranked by fold increase) and transmission route.

| Baltimore classification |  | Family |  |  |
| --- | --- | --- | --- | --- |
|  | Predicted host range<br>(~fold increase) |  |  | Predicted host range<br>(~fold increase) |
| **Group I (dsDNA)** | 8.63 [3, 30.43] (~2.59<br>[~1.16, ~6.94]) | **Bornaviridae** | V | 71 [15.5, 293.25] (~9.51<br>[~2.08, ~42.22]) |
| **Group II (ssDNA)** | 5.47 [2.19, 24.88] (~2.04<br>[~1.07, ~6.56]) | **Orthomyxoviridae** | V | 60.25 [15, 196.5] (~7.76<br>[~1.62, ~27.19]) |
| **Group III (dsRNA)** | 27.15 [7.96, 93.11] (~4.04<br>[~1.41, ~11.94]) | **Rhabdoviridae** | V | 52.8 [23.68, 149.09] (~7.33<br>[~1.81, ~24.03]) |
| **Group IV ((+)ssRNA)** | 17.64 [5.34, 65.29] (~3.49<br>[~1.26, ~10.9]) | **Hepeviridae** | IV | 70.67 [25.33, 220] (~6.67<br>[~2.49, ~15.54]) |
| **Group V ((-)ssRNA)** | 24.91 [8.36, 100.53] (~4.44<br>[~1.39, ~18.08]) | **Filoviridae** | V | 31.75 [7, 155.62] (~5.77<br>[~1.3, ~25.37]) |
| **Group VI (ssRNA-RT)** | 26.68 [10.26, 94.58] (~4.99<br>[~1.54, ~15.36]) | **Togaviridae** | IV | 48.5 [12.45, 161.65] (~5.71<br>[~1.52, ~16.95]) |
| **Group V (dsDNA-RT)** | 19.29 [7.29, 109.43] (~2.53<br>[~1.35, ~14.55]) | **Flaviviridae** | IV | 40.59 [11.26, 131.77]<br>(~5.09 [~1.37, ~16.14]) |
|  |  | **Retroviridae** | VI | 26.68 [10.26, 94.58] (~4.99<br>[~1.54, ~15.36]) |
| **Transmission route** |  | **Coronaviridae** | IV | 22.86 [6.23, 94.89] (~4.81<br>[~1.44, ~17.85]) |
| **Direct** | 14.67 [5.07, 55] (~3.29<br>[~1.25, ~10.49]) | **Poxviridae** | I | 23.22 [7.76, 88.76] (~4.77<br>[~1.39, ~15.74]) |
| **Direct sexual** | 18.26 [6.05, 60.05] (~3.18<br>[~1.27, ~9.17]) | **Reoviridae** | III | 32.5 [9.39, 111.21] (~4.56<br>[~1.49, ~13.71]) |
| **Direct vertical** | 20.26 [6.81, 68.79] (~3.44<br>[~1.31, ~10.48]) | **Paramyxoviridae** | V | 26.28 [9.39, 98.79] (~4.46<br>[~1.61, ~14.4]) |
| **Indirect** | 20.02 [7.35, 71.38] (~3.41<br>[~1.27, ~11.47]) | **Phenuiviridae** | V | 26.94 [6.35, 124.18] (~4.25<br>[~1.23, ~20.34]) |
| **Ingestion** | 10.7 [4.11, 39.46] (~2.52<br>[~1.15, ~7.69]) | **Peribunyaviridae** | V | 20.09 [5.85, 90.45] (~4.15<br>[~1.39, ~19.03]) |
| **Inhalation** | 14.53 [4.6, 59.87] (~3.29<br>[~1.24, ~11.73]) | **Hantaviridae** | V | 15.61 [4.83, 77.59] (~3.61<br>[~1.23, ~17.13]) |
| **Environmental** | 20.32 [6.25, 82.58] (~3.81<br>[~1.29, ~14.12]) | **Picornaviridae** | IV | 13.62 [4.45, 52.93] (~3.55<br>[~1.32, ~10.7]) |
| **Vector** | 30.1 [8.38, 117.44] (~4.73<br>[~1.42, ~18.26]) | **Pneumoviridae** | V | 28.89 [8.67, 107.56] (~3.47<br>[~1.18, ~12.78]) |

### Mammalian reservoirs of viruses

#### Relative importance of mammalian features

our suggested separation of perspectives allowed us to calculate relative importance of mammalian traits (Table 2) to our viral models. We were also able to capture variations in how these features contribute to our viral models at various levels (e.g. Baltimore classification, or transmission route) as highlighted in Figure 3 (panel A). Our results show that distances to known hosts of viruses were the top predictor of associations between the focal virus and our terrestrial mammals. The breakdown was: 1) mean phylogenetic distance - all viruses = 98.75% [93.01%, 100%], DNA = 99.48% [96.03%, 100%], RNA = [91.93%, 100%]; 2) mean ecological distance all viruses = 94.39% [71.86%, 100%], DNA = 96.36% [80.99%, 100%], RNA = [69.48%, 100%]. In addition, life-history traits significantly improved our models, in particular: longevity (all viruses = 60.9% [12.12%, 98.88%], DNA = 68.03% [11.22%, 99.69%], RNA = [13.55%, 96.37%]); body mass (all viruses = 62.92% [5.4%, 97.65%], DNA = 72.75% [18.49%, 100%], RNA = 57.45% [4.32%, 95.5%]); and reproductive traits (all viruses = 53.37% [5.67%, 95.99%]%, DNA = 59.46% [8.27%, 99.32%], RNA = 50.17% [4.85%, 92.17%]).

#### Wild and semi-domesticated mammalian reservoirs of viruses

our framework indicated ∼4.28 -fold increase [∼1.2, ∼14.64] of the number of virus species in wild and or semi-domesticated mammalian hosts (16.86 [4.95, 68.5] viruses on average per mammalian species). These results indicated an average of 13.45 [1.73, 65.04] unobserved virus species for each wild or semi-domesticated mammalian host (known viruses that are yet to be associated with these mammals). Our framework highlighted differences in the number of viruses predicted per order (Table 4). Figure 3 illustrates the predicted number of viruses in wild or semi-domesticated mammal by mammalian host range (panel B), and the top 18 virus genera (per number of host-virus associations) in selected orders (panel C).

**Table 4.** Predicted number of viruses per top 15 orders by fold increase in number of viruses predicted in wild or semi-domesticated mammalian hosts. Results are ordered by descended fold increase.

| Order | Fold increase | Virus range | New viruses |
| --- | --- | --- | --- |
| Tylopoda | ~12 [~1.75, ~30.45] | 31.5 [4, 125] | 29 [1.5, 122.5] |
| Ruminantia | ~9.12 [~1.69, ~23.53] | 41.45 [10.92, 126.55] | 36 [5.61, 121.07] |
| Primates | ~7.12 [~1.48, ~20.88] | 34.46 [10.24, 114.62] | 27.77 [3.86, 107.89] |
| Suina | ~7.12 [~0.88, ~28.01] | 21.83 [3.33, 112.83] | 18.92 [1.33, 109.83] |
| Perissodactyla | ~5.74 [~1.67, ~17.78] | 25.18 [8, 97.09] | 20.64 [3.73, 92.55] |
| Cingulata | ~5.67 [~1.67, ~27] | 17 [5, 107] | 14 [2, 104] |
| Lagomorpha | ~5.27 [~1.9, ~16.43] | 18 [5.25, 74.83] | 15.08 [2.42, 71.92] |
| Rodentia | ~4.79 [~1.18, ~18.72] | 15.22 [3.54, 74.84] | 12.65 [1.26, 72.22] |
| Carnivora | ~3.91 [~1.23, ~14.84] | 18.3 [5.99, 78.38] | 14.11 [1.94, 74.15] |
| Hippopotamidae | ~3 [~1, ~10.5] | 3 [1, 20] | 2 [0, 19] |
| Chiroptera | ~2.79 [~1.08, ~9.5] | 9.51 [3.11, 41.66] | 7.11 [0.84, 39.24] |
| Scandentia | ~2.15 [~0.73, ~10.34] | 11 [4.33, 55.33] | 5.67 [1, 50] |
| Didelphimorphia | ~2.13 [~0.93, ~8.57] | 4.6 [1.6, 27] | 3 [0, 25.2] |
| Eulipotyphla | ~2.04 [~0.86, ~13.78] | 7.14 [2.45, 55] | 4.73 [0.36, 52.45] |
| Pilosa | ~1.84 [~0.6, ~17.98] | 7.8 [3.8, 68.4] | 3.4 [0, 63.8] |

## DISCUSSION

Overall, we predict a 5.35-fold increase in associations between wild and semi-domesticated mammalian hosts and zoonotic viruses (found in humans, excluding rabies virus). Similarly, our results indicate a 5.2-fold increase between wild and semi-domesticated mammalian hosts and viruses of economically important domestic species (e.g. livestock and pets).

Figure 2C illustrates the increase in number of predicted associations per mammalian order. For bats and rodents which have been associated with recent outbreaks of emerging viruses such as coronaviruses^43^ and hantaviruses^44^, there are 5.55-fold and 5.45-fold increases respectively (supplementary table S15). The fold increases are higher for zoonotic viruses and viruses observed in economically important domestic species, where for bats we predict a 7.42-fold and an 8.29-fold increase respectively, and for rodents we predict a 7.42-fold and a 7.7-fold increase, respectively. The increase in associations indicated a non-uniform knowledge-gap across mammalian virus reservoirs. For bats the largest fold increase was in group III viruses with an 8.72-fold-increase, whereas in rodents the highest fold increase was in group V viruses - a 6.23-fold-increase.

Figure 3C illustrates the top viral genera (per number of predicted associations) in important mammalian reservoirs including bats and rodents. The largest significant fold increases in bats were with the group V Lyssaviruses (excluding rabies virus), a family of viruses causing an array of medically and veterinary important rabies-like diseases in a wide range of mammals^45,46^, with a 10.4-fold increase in the number of predicted associations (figure 3C, table S15). Group V Bornaviruses, which cause a range of encephalitic diseases in mammals including the fatal Borna disease^47^ (sad horse disease) common in horses and other domesticated animals, had a 23 and a 12-fold increase in associations in bats and rodents, respectively. Finally, group III Rotaviruses had an 8.11-fold increase in bats – rotaviruses are the most common cause of diarrhoeal diseases in children and are of particular concern in developing countries^48^.

Analogous to bats and rodents being important reservoirs of zoonotic viruses, wild ruminants are key in maintaining and circulating of viruses affecting ruminant livestock^49^. Our model highlights this knowledge-gap by predicting a 7.77-fold increase in number of associations between wild and semi-domesticated ruminants and known viruses (figure 2C, table S14); and a 10.11-fold increase in associations between these ruminants and observed zoonotic viruses. Furthermore, our model predicted significant increase in mammalian host range of significant livestock viruses including: a 7.45-fold increase in range of Venezuelan equine encephalitis virus (Group IV, Togaviridae); a 5.33-fold increase in range of Schmallenberg orthobunyavirus (Group V, Peribunyaviridae); and a 2.96-fold increase in range of bluetongue virus (Group III, Reoviridae).

These results show that our approach is able to highlight significant numbers of missing associations of medically- and veterinary-important viruses and their reservoirs. Such information can be used to better understand the risk to people and livestock from these reservoirs and guide policy to take appropriate precaution to mitigate infection.

There are a number of reasons for which virus-mammal associations may have been disproportionately under-described, which can be categorised as follows. *Public health, food security and economically driven research biases*: Most of our current knowledge of infectious agents, including viruses, is centred upon humans. Second to humans (37.1% of captured mammalian research effort), agricultural and companion animals tend to receive significantly more research effort (∼15% of captured mammalian research effort). Examples include the well-studied microbiome of domestic cats (*Felis catus*, 57 known virus species) compared with the understudied microbiome of wild felines (e.g. *Felis silvestris*, 13 known virus species – these expanded to 51 viruses using our framework). Linked to this is wealthier countries producing a larger research volume, and hence interactions common within or of importance to such countries are more likely to be described.

### Practical limitations

infectious agents of endangered and rare mammalian species, and mammalian species found predominantly in remote regions are less likely to be characterised due to difficulties in sampling these mammals in their natural habitats. The same applies to viruses with restrictive geographical range, or those that are less common in mammals (e.g. avian pathogens). Our framework was able to expand the host range of rare viruses, found in one or two mammals (N=1,450) from 1,619 to 4,174 hosts (∼ 2.16 average increase per rare virus). It also increased the virus range of rarely studied mammals (n=954) from 1,150 to 4,318 viruses (∼3.21 average fold increase per host).

### Biological reasons

virus-mammal associations which produce more visible or marked effects are more likely to be studied. Examples include fertility or physically observable interactions being over-studied, whilst potentially important asymptomatic interactions, or interactions where a cross-immunity from related viruses masks observable symptoms may remain unnoticed and hence understudied. Furthermore, co-evolution between virus and primary host often results in a less severe phenotype, whilst the same virus in an incidental host may result in more marked and hence more studied disease. Studied examples include Ebola virus presenting minimal symptoms in bats but extreme symptoms in humans; analogous interactions where the former host may have been unobserved are likely to be plentiful. For example, our framework indicated that 19 additional species of bats could be carrier of Ebola virus in addition to 10 known species.

The novelty of our approach lies in the separation of perspectives - by isolating the viral, mammalian and network perspectives we were able to further our understanding of mammalian reservoirs of known viruses as follows: 1) our novel divide-and-conquer approach explored the explanatory power, by means of variable importance, of a comprehensive set of mammalian and viral traits. Uniquely, we incorporated geospatial features extrapolated from an extensive collection of global data on climate, environmental, agricultural, and mammalian diversity variables. 2) We consolidated these viral and mammalian traits with network topological features, expressed in terms of *potential motifs*. By counting potential motifs - in which an unknown virus-mammal association (link) may feature - we were able to quantify the topology of our network and incorporate this topology into the prediction process in an explainable, measurable, and extendible way. 3) Our voting approach, despite being more conservative than its components (our 3 perspectives, supplementary results 2-4), was able to bridge a significant gap in our knowledge of reservoirs of mammalian viruses (18,920 associations between wild and semi-domesticated mammalian species and known viruses).

There remains, however, key areas for further improvement. Prediction of novel viruses and their potential threat to humans, livestock and wildlife is an increasingly important and active research area. Where an established virus is increasing its range beyond the native region (e.g. due to climatic or demographic factors), then our framework provides powerful means to assess potential hosts it has yet to come into contact with. However, for completely novel (e.g. SARS-CoV-2) or never-studied viruses our approach cannot predict potential associations. Future work may be able to enhance the predictive power of our approach by incorporating more diverse viral traits, particularly in terms of detailed genetics as proposed recently^7^ and in terms of geographical distribution and associated features of the virus as highlighted in previous work^50,51^. Integration of predictors of the host-virus interactions such as existence of particular viral receptors in host cells would also greatly benefit our models and create a fourth perspective that could be added into the voting process. Finally, further separation of perspectives could also be achieved by incorporating arthropod vectors or intermediate hosts, in effect creating a tripartite network for which our framework could be extended. Additional perspectives could potentially be added for different classes of hosts, particularly birds, many species of which are also reservoirs or amplifying hosts of important viruses (e.g. West Nile and Japanese encephalitis viruses). Conversely it will be interesting to further expand the pathogens included in our models to incorporate bacterial, protozoal, helminth and fungal pathogens.

## METHODS

### Virus-host species associations and bipartite network formulation

We extracted species-level virus-mammal associations from the Enhanced Infectious Diseases Database^8^ – EID2. We recursively aggregated virus-mammal associations – a mammal that was found to host a strain or subspecies of virus was considered a host of the corresponding virus species. We further checked these species-level associations for accuracy and to eliminate laboratory-produced associations and spurious instances. This resulted in 6,331 associations between 1,896 viruses and 1,436 terrestrial mammals. We transformed these associations into a bipartite network in which nodes represent either virus or mammal species, and links indicate associations between mammalian and viral species (Supplementary Note 1).

### Multi-perspective framework to predict unknown virus-mammal associations

Our bipartite virus-mammal network is sparsely connected – roughly 0.23% of potential associations are documented in EID2, despite it being the most comprehensive resource of its kind. This sparsity is more evident in wild and semi-domesticated species where only 0.182% of potential associations are observed. We treated the problem of bridging this gap in our knowledge of virus-mammal associations as a supervised classification problem of links in the bipartite network. In other words, we aimed to predict unknown associations between known viruses and their mammalian hosts based on our knowledge to date of these species. Each possible virus-mammal association is predicted three times as follows.

#### 1 From the mammalian perspective

For each mammal in our network, given a set of features (predictors) comprising viral traits (e.g. genome, transmission routes) – table 1, what is the probability of an association forming between this mammal and each of the 1,896 virus species?

#### 2 From the viral perspective

For each virus species found in our network, given a set of features (predictors) encompassing mammalian phylogeny, ecology and geographical distribution – table 2, what is the probability of an association forming between this virus and each of 1,436 terrestrial mammals?

#### 3 Form the network perspective

Given a set of topological features representing the bipartite network expressing most of our knowledge to date of virus-mammal associations, what is the probability of an association forming between any virus and any mammal in our dataset (N=1,896 x 1,436 = 2,722,656 possible associations)?

Our framework trained and selected a set of supervised classifiers in each of the above perspectives as discussed below. It then consolidated the results of the best performing classifiers using voting whereby an unknown (potential or unobserved/undocumented) association was selected if it was predicted by at least two of the three perspectives.

Our framework is flexible, both in terms of machine-learning algorithms selected, classifiers trained, and features engineered for each perspective. It avoids overfitting as it approaches the problem from various perspectives, and effectively consolidates ensembles of classifiers trained on subsets of the underlying data. In addition, no constituent model of our framework has been trained with all available data at any time. Finally, our framework enables the incorporation of hosts where only one virus has been detected to date (via perspectives 2 and 3), and viruses where only one host has been discovered (via perspectives 1 and 3).

### The network perspective - Topologically derived network features of virus-mammal associations

In contrast with our mammalian and viral perspectives, the network linking known viruses with their mammalian hosts presents a global view of how these viruses are shared amongst mammalian hosts. Here we capture the topology of this bipartite network by means of counts of *potential motifs*^16^. Figure 4 illustrates 3, 4, and 5 node motifs which might appear in a bipartite virus-mammal association network. These motifs capture important indirect pathways between viruses and their mammalian hosts. These pathways vary from simple generalisations capturing whether a virus has wide range of hosts or not (m3.1, m4.1, and m5.1), or if the mammal is exposed to many viruses (m32, m46, m520), to more complex pathways (e.g. two host species sharing 80% of their viruses with each other; three viruses sharing 50% of their hosts with each other). These pathways might indicate if an unknown association is more likely to exist in nature or not, and are only capturable, and most importantly quantifiable, at the global level as encapsulated by our network perspective.

**Figure 4.**
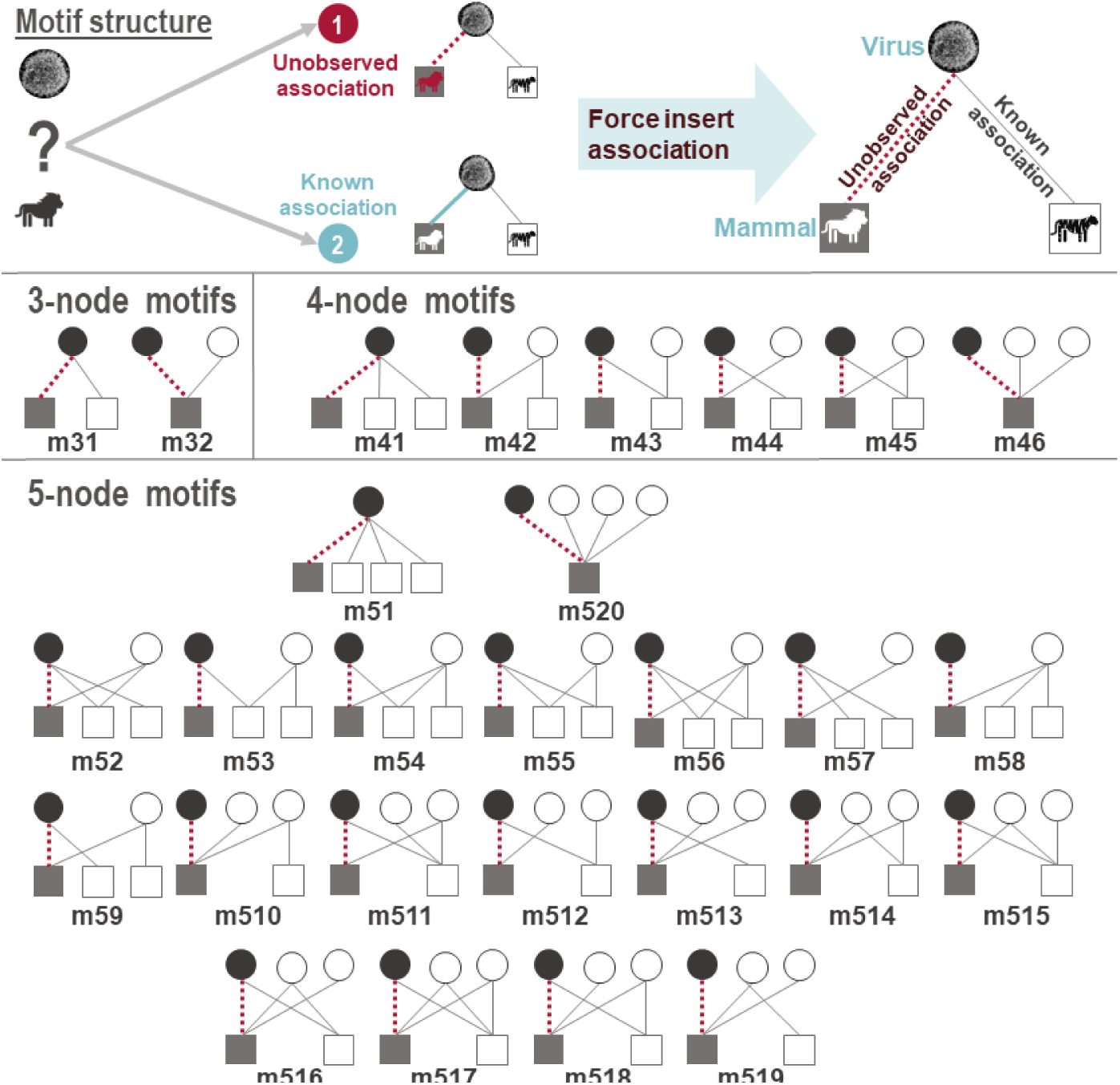
3, 4 and 5 node *potential* motifs in our virus-host bipartite network. Circles represent virus species and square represents mammal species. Dashed red lines represent known or unknown (i.e. potential/missing, unobserved/undocumented, or negative) virus-mammal associations. Grey lines illustrate existing associations in our network of 1,896 virus species and 1,436 terrestrial mammals. Associations between viruses and mammals have two states: known (documented in our sources) or unknown. Unknown associations represent the gaps in our knowledge, they could exist in nature but are undocumented or can exist in the future. Therefore, we force insert each possible virus-mammal association (N=2,722,656) prior to counting motifs (hence termed potential motifs).

Capturing these indirect pathways, by means of motifs, enable us to apply supervised machine learning algorithms to make predictions directly from the network structure, which is not captured by the other two perspectives. Motifs are usually associated with specific frequency thresholds^18^. However, here we follow previous work^16^ in removing this restriction. We simply count the number of occurrences of the motifs outlined in figure 4, as discussed below, and then let the machine learning algorithms detect which motifs are particularly important to the problem of predicting links in our network.

In order to incorporate these motifs as features from which a supervised classifier can learn, we applied the following approach. For each virus-host association (link), whether known or unknown, we counted the number of instances of each of our motifs (as categorised in figure 4) in which the association might feature (presented as dashed red line in figure 4). In other words, for each virus-host association we “*inserted”* the corresponding link into our network and counted all potential motifs in which this link might feature if it actually existed^16^. This enabled us to create a training set of all potential host-virus associations of our 1,896 viruses and 1,436 mammals and the counts of their potential motifs. We then trained a number of machine-learning algorithms with this dataset as detailed in following subsections. **Research effort:** We incorporated research effort on mammal and virus species into our network perspective models. We calculated research effort as the total number of sequences and publications of each species as indexed by EID2^8^.

### Multi-perspective prediction pipeline of unknown virus-mammal associations

As highlighted above, our framework comprised three perspectives: mammalian, viral and network. Each of these perspectives trained a set models with different features (tables 1 and 2, and figure 4 respectively), and hence required its own pipeline as described below (Supplementary Note 5).

#### Mammalian and viral perspectives

##### Class balancing

On average each virus in our dataset affected 3.45 mammals (∼0.24%), and each mammalian host was affected by 4.41 viruses (∼0.24%). This presented an imbalance in our data, whereby a small percentage of instances are actualised. We dealt with this issue in two ways: first we excluded any virus (N=1,281) which was found in only one mammal species from our virus models pipeline (viral perspective), and we excluded any mammal (N=758) which is only affected by one virus from our mammal models pipeline (mammalian perspective). Second, we deployed SMOTE - Synthetic Minority Over-sampling Technique^52,53^ to rebalance the classes prior to training each of our viral (N=8×556) and mammalian (N=8×699) models.

##### Classification algorithms

For each mammal and each virus selected above we trained 8 classification algorithms (supplementary table S4): Model Averaged Neural Network (avNNet), Stochastic Gradient Boosting (GBM), Random Forest, eXtreme Gradient Boosting (XGBoost), Support Vector Machines with radial basis kernel and class weights (SVM-RW), Linear SVM with Class Weights (SVM-LW), SVM with Polynomial Kernel (SVM-P), and Naive Bayes. We selected these classifiers due to their robustness, scalability, availability, and over-all performance^54,55^. All models were trained and tested via *caret* R package^56^.

##### Training & tuning

each of the above models was trained with 10-fold cross validation (Supplementary Note 5). In order to allow us to incorporate uncertainty arising from variations in SMOTE resampling technique and resulting training sets, and to generate empirical confidence intervals we repeated the above method 50 times for each virus or mammal model.

##### Model selection strategy

We computed three performance metrics based on the median predicted probability across each set of replicate models: AUC, true skills statistics (TSS) and F1-score (supplementary table S5). The best performing classifier per each virus or mammal, across all measures, was included in our multi-perspective final model (Supplementary Note 5, Supplementary Results 2 and 3).

#### Network perspective

##### Class balancing

Our bipartite virus-mammal network is sparsely connected with 6,331 documented associations out of 2,722,656 possible associations (0.233%). Due to this network connecting all input mammal and virus species we implemented strict under-sampling: We generated balanced training sets by drawing 1000 known associations and 1000 unknown (negative) associations at random.

##### Training & validation

We trained same selection of algorithms as above with balanced sets (2000 instances each) using 10-fold cross validation to optimise AUC. We repeated this process 100 times to generate a bragging (median) ensemble of predictions (derived as probabilities) of its replicate models. Performance metrics were calculated for each ensemble and the best overall ensemble was incorporated into our final model (SVM-RW - Supplementary Results 4).

#### Performance assessment

We trained constituent models of each perspectives with a stratified random training set comprising 85% of all data. The processes described above were repeated with training set only, and performance was measured against the held-out test set (15% of all data). Performance metrics obtained through this assessment were reported in our results and supplementary results sections. In addition, we performed an additional test to assess the ability of our model to predict systematically removed virus-mammal associations (Supplementary Results 1).

#### Variable importance

we calculated relative importance (influence or contribution) of viral and mammalian features (listed in tables 1 and 2, respectively) to each model in our mammalian and viral perspectives. Due to the selection strategy implemented, we computed the importance of these features via model-independent filter approach via a ROC curve analysis conducted on each predictor (as implemented in the *caret* package^56^).

## Supporting information

Supplementary Notes and Results

## ACKNOWLEDGMENTS

MW acknowledges support from BBSRC and MRC for the National Productivity Investment Fund (NPIF) fellowship (MR/R024898/1). Establishment of the EID2 database was funded by a UK Research Council Grant (NE/G002827/1) to MB, as part of an ERANET Environmental Health award to MB; subsequently, it has been further developed and maintained by BBSRC Tools and Resources Development Fund awards (BB/K003798/1; BB/N02320X/1) to MB, and the National Institute for Health Research Health Protection Research Unit (NIHR HPRU) in Emerging and Zoonotic Infections at the University of Liverpool in partnership with Public Health England and Liverpool School of Tropical Medicine.

