## Supplementary Notes and Results for "Divide and conquer – machine-learning integrates mammalian, viral, and network traits to predict unknown virus-mammal associations"

### Electronic Supplementary Materials

**Maya Wardeh*, Marcus SC Blagrove, Kieran J. Sharkey, Matthew Baylis**

#### Supplementary Note 1 – Virus-mammal interactions data

**Mammal-virus associations:** Species-level virus-mammal associations were extracted from the Enhanced Infectious Diseases Database[1] – EID2. EID2 contains 4,799 species of mammals and 9,605 species of viruses. EID2 extracts information on pathogens, their hosts and locations from two sources: 1) meta-data accompanying nucleotide sequences (hereafter sequences) published in Genbank[2,3]; and 2) titles and abstracts (hereafter TIABs) of publications indexed in the PubMed database[4]. To date, EID2 has extracted information from 71,397,061 sequences (and processed 100M+ sequences), and 8,643,203 TIABs. Here, we recursively aggregated virus-mammal associations – a mammal that was found to host a strain or subspecies of virus was considered a host of the corresponding virus species (and vice versa). We further checked these species level associations for accuracy and to eliminate laboratory-produced results. This resulted in 6,331 associations between 1,896 viruses and 1,436 terrestrial mammals.

**Virus taxonomy:** For the purposes of taxonomic classification (Table S1) of viruses we obtained the most recent taxonomic tree of our viruses from the NCBI taxonomy database [5] (accessed 09/2019).

| | Group | N | H | Description |
|---|---|---|---|---|
| **DNA** | Group I | 726 | 491 | dsDNA viruses - double-stranded DNA viruses (e.g. herpesviruses) |
| | Group II | 273 | 160 | ssDNA viruses - single-stranded DNA viruses (e.g. circoviruses) |
| **RNA** | Group III | 46 | 156 | dsRNA viruses - double-stranded RNA viruses (e.g. rotaviruses) |
| | Group IV | 427 | 511 | (+)ssRNA viruses - positive-sense single-stranded RNA viruses (e.g. Zika virus, Yellow fever virus) |
| | Group V | 360 | 1051 | (-)ssRNA viruses - negative-sense single-stranded RNA viruses (e.g. Ebola virus, Influenza A virus) |
| **Retro-transcribing** | Group VI | 57 | 182 | ssRNA-RT viruses - (+ strand or sense) RNA with DNA intermediate in life-cycle (e.g. retroviruses such as HIV1) |
| | Group VII | 7 | 41 | dsDNA-RT viruses - DNA with RNA intermediate in life-cycle (e.g. Hepatitis B virus) |

**Table S1 – Baltimore classification of mammalian viruses included in this study.** N indicates number of virus species in each group. H indicates number of mammal species for which associations with viruses in each group were found in EID2.

**Domestication level of mammals:** We classified the domestication status of our mammalian hosts into four levels: wild mammals (N=1326), semi-domesticated mammals (N=81), domesticated mammals (N=28), and human (N=1). Table S2 lists our domesticated mammals. Our semi-domesticated mammals group included N=29 of each Carnivora and Ruminantia, 13 Rodentia, 3 Perissodactyla, 2 of each Proboscidea and Tylopoda, and one of each Diprotodontia, Eulipotyphla and Suina.

| | N | Species |
|---|---|---|
| **Artiodactyla** | 16 | *Bison bonasus, Bos frontalis, Bos grunniens, Bos indicus, Bos javanicus, Bos taurus, Bos mutus, Bubalus bubalis, Bubalus carabanensis, Capra hircus, Ovis aries, Camelus bactrianus, Camelus dromedaries, Lama glama, Lama pacos, Sus scrofa* |
| **Carnivora** | 3 | *Canis lupus familiaris, Felis catus and Vulpes vulpes.* |
| **Perissodactyla** | 3 | *Equus asinus, Equus caballu and Equus asinus x caballus* |
| **Rodentia** | 5 | *Cavia porcellus, Mesocricetus auratus, Mus musculus, Rattus norvegicus and Rattus rattus.* |
| **Lagomorpha** | 1 | *Oryctolagus cuniculus* |

**Table S2 – Domesticated mammals included in our analyses.**

#### Supplementary Note 2 – Viral Feature Space (Viral traits & features)

**Virus genome and capsid:** We classified the genome of each virus as RNA (binary factor, no= DNA); retro-transcribing (binary factor); negative-sense (binary factor, yes = negative-sense; and positive-sense (binary factor, yes = positive -sense). RNA viruses adapt faster [6], and are generally more fragile (cannot survive as long outside of the cell). Retroviruses are generally very conserved[7], and have to enter the nucleus[8] and insert into the genome, these additional steps may require specificity and limit range. Sense affects replication cycle and range of host enzymes needed.

With regards to the genome architecture and organisation we checked if the virus has circular or linear genome (binary factor, no=linear), as this attribute affects replication and translation. Rolling circle replication and translation are common with circular genomes, negating the need to re-enlist host enzymes[9], therefore possibly affecting host range as less specificity may be required. In addition, we noted if the virus was monopartite (has a single nucleic acid molecule protected in a shell made of proteins) or segmented (divided into two or more nucleic acid segment) (binary factor, no=monopartite). Segmented viruses can undergo recombination if two strains of the same virus infect a cell (e.g., influenza hemagglutinin & neuraminidase recombination[10]). This in turn can lead to host range changes of segments of the genome. In practice, there exists a third class of viral architecture - multipartite viruses. These viruses have their genome divided into two or more nucleic acid segment (similarly to segment viruses), but these segments are each packaged into separate virus particles. We ignored multipartite viruses in this study due to them being very rare and poorly understood[11].

Regarding capsids, we indicated if the virus is enveloped or not (binary factor). Envelopes are usually derived from the host cell membrane; this can help them avoid host immune system. They may limit range by providing antigens for other immune systems. The envelopes are very sensitive to the external environment, and enveloped viruses often require to be directly transferred between hosts; finally, because the envelope is made from the current host's cell membrane, it will change upon infection of a new host, making the virus rapidly adaptable[12].

GC-content (guanine-cytosine content), the percentage of a nucleotide sequence that is made up of either guanine or cytosine bases of each virus, and the average genome size (in bases) were also obtained for our viruses. GC hydrogen bonding is stronger than AT/U, high GC content usually leads to higher thermo-stability of the genome[13], including single stranded genomes which often self-anneal. This may affect longevity outside of hosts, and replication inside of ectothermic hosts. Genome size is indicative of many aspects of the virus such as complexity, DNA/RNA, and replication type.

**Virus replication, release, and cell entry:** we collated information on the replication site of the virus. We expressed these data as a binary factor indicating if the virus replicates in the cytoplasm (versus nucleus replication). Replication site is linked to RNA/DNA genome – if a virus has a DNA stage it must replicate in the nucleus. This creates extra barriers to overcome for entry to the nucleus and may restrict host range. We classified the release of the virus into three broad categories: budding, lysis, or other. The mechanism of release affects aspects the rate of virus production, cell life-span and means of presentation to the immune system[14], each of these aspects could influence the virus host range. Finally, we recognised that availability of receptors influences potential host range, therefore we broadly categorised the mechanism of virus cell entry into 4 categories: cell-receptor endocytosis, clathrin-mediated endocytosis, glycoprotein-mediated or other.

**Transmission routes:** We categorised major transmission routes of our viruses as follows: **direct transmission** – by direct contact, via skin, broken sink or droplets (unique viruses $N_v$=1,168, mammalian hosts $N_m$=1305, associations=4,389); **sexual transmission** ($N_v$=273, $N_m$=405, associations=1,154); **vertical transmission** – mother to child, or via breast-milk contact ($N_v$ = 252, $N_m$=445, associations =1,161); **indirect transmission** – e.g. via secretions/excretions/tissues ($N_v$ =

602, $N_m$=1260, associations =3,224); **ingestion** – including water ingestion and faecal-oral routes ($N_v$ = 848, $N_m$=1,013, associations =2,792); **inhalation** – via droplet or airborne particles or dust ($N_v$ = 641, $N_m$=612, associations =2,029); **environmental** – through fomite, contact with environment ($N_v$ = 372, $N_m$=633, associations =1,596); and vector – strictly via arthropod vector such as ticks or mosquitoes ($N_v$ = 194, $N_m$=377, associations =1,019).

We adopted a three-fold strategy to capture major transmission routes of our viruses as follows:

1. Title and abstract (TIABs) mining: we utilised EID2 to extract TIABs of PubMed papers linked to single virus species (i.e. excluding TIABs with multiple viruses). These TIABs were subsequently classified via keyword matching into the transmission routes described above. The TIABs were further checked manually to ensure correctness and to remove transmission routes outside mammalian species (e.g. we removed instances of vertical transmission within arthropod vectors).
2. Manual extractions from online-sources: we manually extracted transmission routes of viruses for which no papers were identified by the previous step, as well as for routes not detected in the TIABs from various sources[15–18].
3. Within genus generalisations: finally, we assigned transmission routes to viruses not captured by the previous two steps, by taking a minimal agreement set of within-genus transmission routes.

For the purposes of our study we utilised a simple multi-label classification of our viruses whereby routes were assigned either 1 or 0 value corresponding to whether the virus was found to be transmitted via the specified route as described above.

#### Supplementary Note 3 – Mammalian Feature Space (mammalian traits & features)

**Phylogeny**: Host phylogenetic distance has been found to drive sharing of pathogens, particularly viruses[20–22], between mammals. We utilised a recent mammalian supertree[23] to calculate pairwise phylogenetic distance between each mammal-mammal pair. We then aggregated these values (mean) between each mammal species and the known hosts of each of our viruses. This measure indicated, per virus, whether a potential host species was *phylogenetically* close to or distant from the viruses preferred host range. Mammalian species for which could not be matched to this phylogeny was dropped from our analyses.

In addition, we calculated the evolutionary distinctiveness for each mammal using fair proportion[24], as implemented in the R package *picante*[25]. Evolutionary distinctiveness quantifies how isolated a species is on its phylogenetic tree[26], and has been shown to correlate negatively with pathogen species richness[27].

**Host traits:** We compiled data on morphological and life-history traits, diet and habitat for our mammal species from online databases and literature[28–33]. We selected the following traits for their known correlation with host-pathogen associations, and their wide availability. *Body mass* (g) represented morphological traits, as it proxies key features of metabolism and adaption to environment. For life-history traits we included: *maximum age* (months), *activity cycle*, and *migration*[29]. We included *gestation period length* (days), *litters per year*, *litter size, weaning age* (days), and *age at sexual maturity* (days), to represent the reproductive characteristics of our mammalian species. Reproductive traits could be viewed as proxies to within-host virus-dynamics and therefore may influence the viruses harboured by the host.

We included geographical area range[32] (in squared km) as species with wider areas might be exposed to more viruses. We incorporated habitat utilisation[32] as multiple binary indicators of whether a species uses one or more of 14 natural and artificial habitats. We hypothesised that mammals utilising similar habitats might come into contact with similar viruses, and this in turn would increase the chances of being infected with these viruses.

We used the proportional use of 10 diet categories[31] to indicate the dietary preferences of mammals. We incorporated these categories as independent variables in our models as we assumed that similar dietary habit might associate with similar viral assemblage.

We included the above listed traits as independent variables in our *virus perspective* models. In addition, we utilised them to quantify the pair-wise ecological distance between each pair of mammals in our study. We based this distance on a generalised form of Gower's distance matrices[34,35]. We incorporated these distances to compute the mean distance between each mammal and known host of each virus (and vice versa). Similarly, to the mean phylogenetic distance listed above, the mean ecological distance indicated, for each virus, whether a potential host species was *ecologically* close to or distant from the viruses preferred host range.

##### Mammalian geospatial features

The geographical distribution of host species influences the pathogens with which it might come into contact. Geography also correlates with other factors such as climate, natural environment, and agricultural practices. Climate has been shown to influence a number of human and domestic mammal pathogens (including viruses)[36–38]. Furthermore, climate indirectly affects certain groups such as vector-borne viruses (e.g. Zika, bluetongue and West Nile viruses) through the direct effect it has on the viability of the associated arthropod vectors[39,40] and their capacity to transmit pathogens. In addition to climate, other geographical factors such as biodiversity (species richness), land cover type, agriculture and farming practices, urbanisation and human population have been found to influence certain categories of host-pathogen associations[41,42].

We obtained and processed species-presence maps of mammalian species[32,43,44]. We supplemented these with grids expressing climate[45], mammalian diversity[46], human population[44], land cover (including urbanisation)[47], agriculture[47,48], and distribution of livestock[43]. This allowed us to generate the following geographical features of mammalian species: 1) we expressed *climate* in two features: mean temperature, and mean precipitation. 2) We quantified the diversity of *natural* land cover type (not directly associated with humans). 3) We quantified *agricultural* (including land-cover associated with humans e.g. managed vegetation) and *farming* practices (expressed in number of domesticated livestock and poultry in the species presence area) as entropies of the associated values in the species range. 3) We computed *urbanisation* as percent of urban land in the species presence area. 4) We summed *human population* in the species presence area. Finally, 6) we computed average *mammalian diversity* in in the species presence area.

**Species-presence maps**: We obtained species-presence maps for majority of our mammalian species from IUCN[32]. We extrapolated livestock (including horses) species-presence maps from most recent global distribution maps [43]. Finally, we inferred presence-maps for three domesticated species - dogs (*Canis lupus familiaris*), cats (*Felis catus)* and guinea pigs (*Cavia porcellus*) from Gridded population of the world maps[44], by assuming they co-exist with humans where there is sufficient human populations (n>100). We used the same gridded population maps to extrapolate human species-presence map (n>0). All our geographical maps manipulation was done in QGIS. We dropped any mammalian species for which no presence maps could be derived from our models trained in viral perspective.

**Geospatial features:** we utilised several sources to extract geo-attributes of our mammalian species. Table S3 lists these sources, the features derived, and reason for inclusion in our analyses. Figure S1 illustrates this process. We derived the following geo-features for our mammalian species:

1. Climate attributes:
    a. Mean temperature: mean of monthly temperatures recorded in the species-presence area, averaged between years: 1900-2010[45].
    b. Mean precipitation: Sum of monthly rainfall (precipitation) recorded in the species-presence area, averaged between years: 1900-2010[45].
2. Natural land cover diversity: calculated as Shannon’s entropy of mean percent of land covered with each of the attributes listed in table S3 under category (Natural) Land-cover.
3. Agriculture and farming diversity: calculated as Shannon’s entropy of mean percent of land utilised for the following categories: managed/cultivated vegetation, regularly flooded vegetation, cropland and pasture, and the sum of each of the livestock, horses and poultry listed in table S3.
4. Human population: human population in the species-presence area calculated by intersecting with a recent gridded human population dataset[44].
5. Urbanisation: mean percent of urban land in the species-presence area.
6. Mammalian diversity: mean mammalian diversity[46] in the species-presence area.

| Category | layers(s)/geo-attributes | Source | Res | Reason |
|---|---|---|---|---|
| **(Natural) Land-cover** | Evergreen/deciduous needle-leaf trees (%) | EarthEnv[47] | 0°0′30″ | Type of land cover has been associated with distribution of various mammals [49]. It potentially increases chances of contact between mammalian reservoirs of different viruses. |
| | Evergreen broad-leaf trees (%) | | | |
| | Deciduous broad-leaf trees (%) | | | |
| | Mixed/other trees (%) | | | |
| | Shrubs (%) | | | |
| | Herbaceous vegetation (%) | | | |
| | Barren land (%) | | | |
| **Agriculture & farming** | Managed/Cultivated Vegetation (%) | EarthEnv[47] | 0°0′30″ | |
| | Regularly flooded vegetation (%) | | | |
| | Cropland (%) | HYDE[50] | 0°5′ | |
| | Pasture (%) | | | |
| | Cattle (head count) | Global distribution data (livestock)[43] | 0.0833° | Livestock farming has been linked to emergence and cross-species transmission of number of viruses (e.g. Nipah virus, influenza viruses)[51]. |
| | Sheep (head count) | | | |
| | Buffalo (head count) | | | |
| | Pigs (head count) | | | |
| | Horses (head count) | | | |
| | Chicken (head count) | | | |
| | Duck (head count) | | | |
| **Human** | Human population | SEDAC[44] | 0°5′ | Urbanisation and human population density have been shown to be drivers of disease emergence and spill-over through wildlife-domestic-human interface [42,52,53]. |
| | Urban land (%) | EarthEnv[47] | 0°0′30″ | |
| **Climate** | Mean temperature | CRUTS3[45] | 0°5′ | Climate has been shown to influence a number of human and domestic-mammals pathogens[36–38]. Furthermore, climate indirectly affects certain groups such as vector-borne viruses (e.g. Zika, Bluetongue and West Nile viruses) through the direct effect it has on the viability of the associated arthropod vectors[39,40]. |
| | Mean precipitation | | | |
| **Mammalian diversity** | Number of different mammalian species in a grid cell. | SEDAC[46] | 0°5′ | Mammalian species present in mammal rich areas might be exposed to diverse viruses[54–57]. |

**Table S3 - List of geographical predictor layers integrated within our framework.**

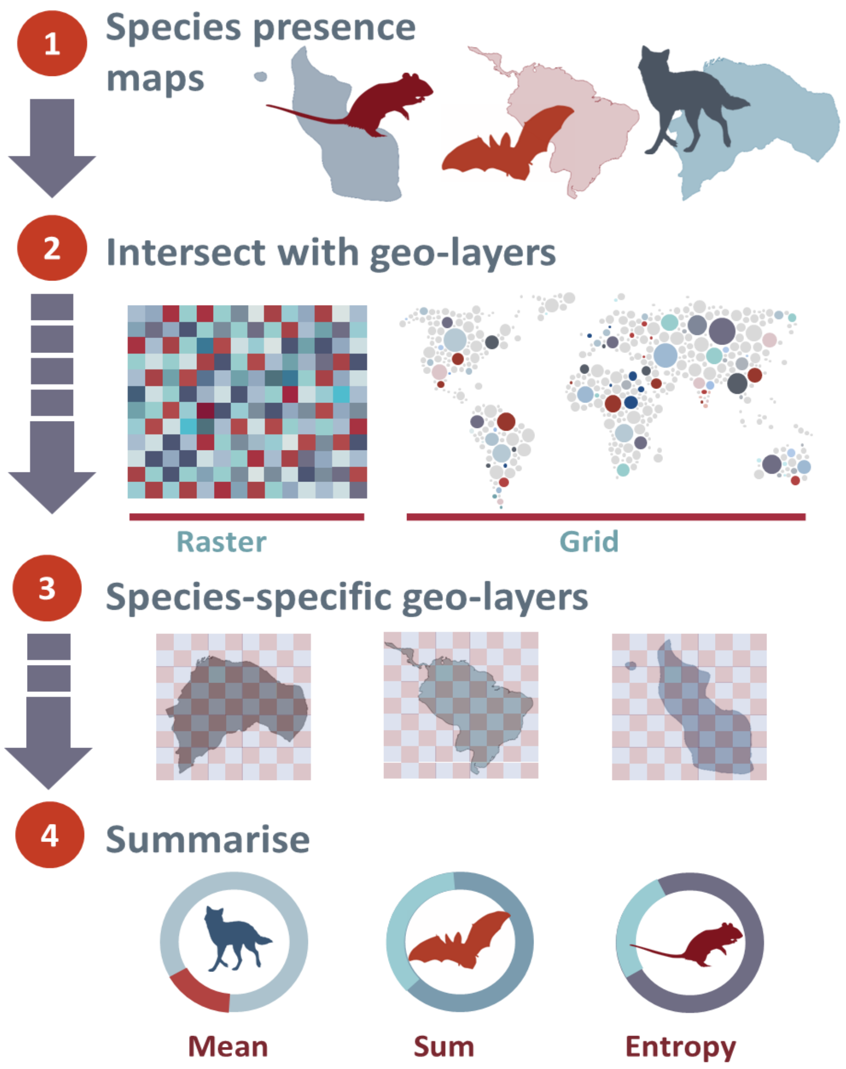


**Figure S1 - Geographical feature extraction**. Mammalian species-presence maps were first extracted from our sources[32,43,44] – step 1; these maps were then intersected with our geo-layers (table S3) – step2. This enabled us to derive values of our geo-attributes for majority of our mammalian species – step3, which we then summarised into the geo-features included in our models – step 4.

#### Supplementary Note 4 – Potential motifs in bipartite networks

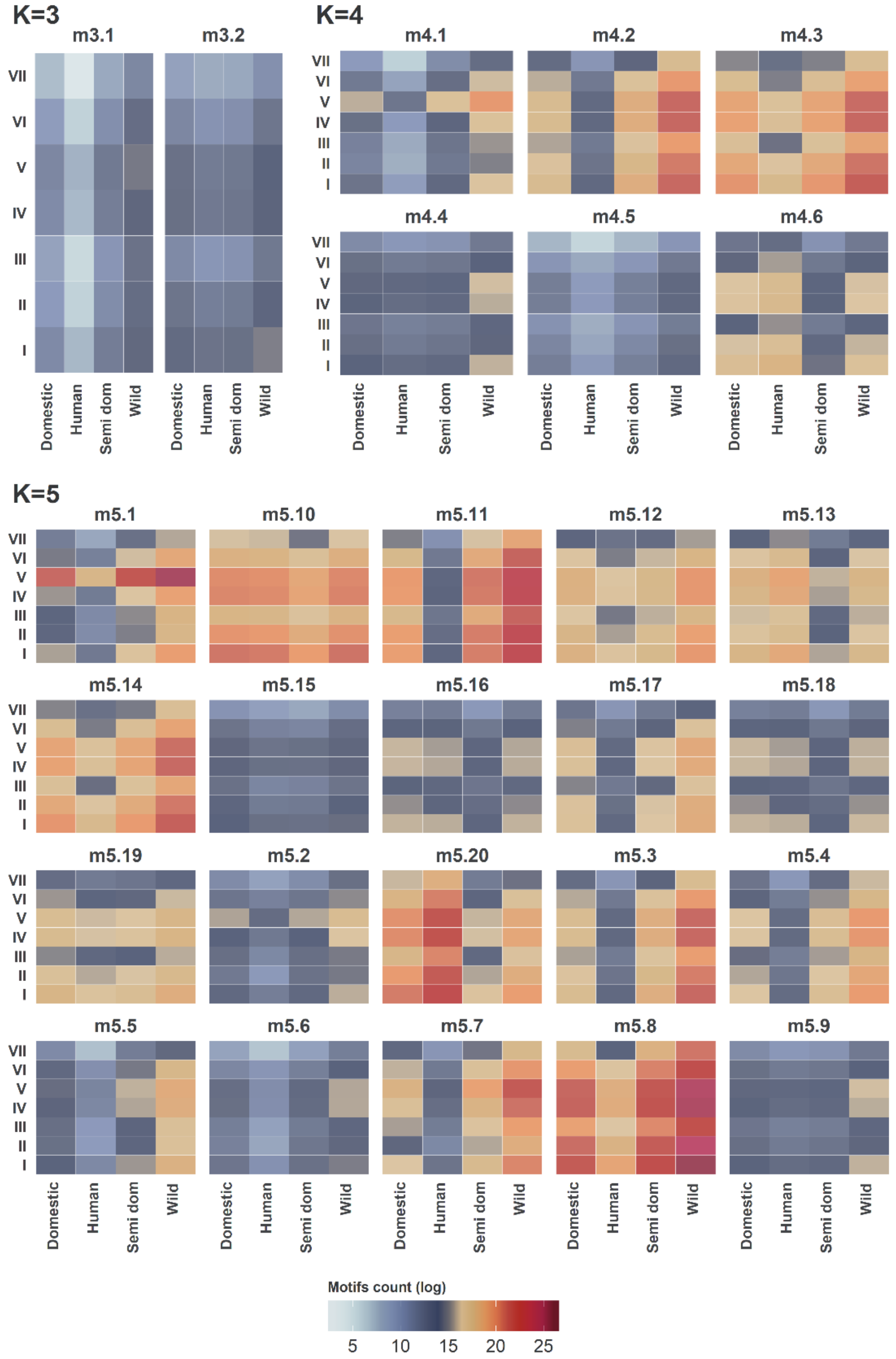


**Figure S2 – Motifs as features of bipartite networks (domestication status).** Heat-maps illustrating distribution of motif-features (counts of potential motifs per each possible edge) in our bipartite network, grouped by domestication status of mammalian hosts and Baltimore classification of the viruses. *The counts are logged to allow for better visualisation.*

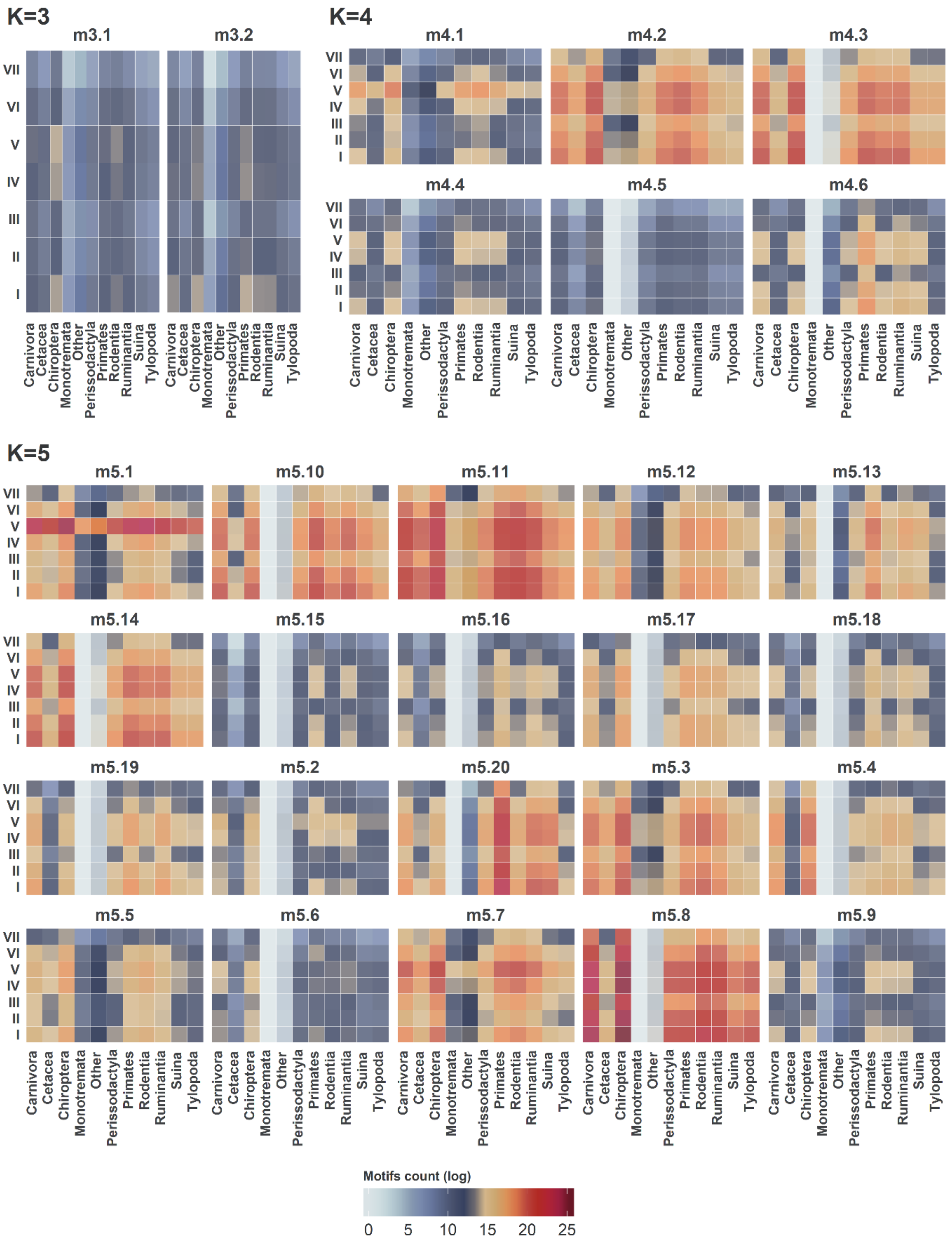


**Figure S3 – Motifs as features of bipartite networks (mammalian order).** Heat-maps illustrating distribution of motif-features (counts of potential motifs per each possible edge) in our bipartite network, grouped by domestication status of mammalian hosts and Baltimore classification of the viruses. *The counts are logged to allow for better visualisation.*

**Supplementary Note 5 – Multi-perspective framework to predict unknown virus-mammal associations**

Machine learning algorithms selection

Table S4 lists the machine learning algorithms (classifiers) included in each of the three perspectives of our framework (N=8). These classifiers offer a varied subset of plethora of classifiers available for experimentation (over 179 classifiers categorised into at least 17 families[58]), and were selected due to their robustness, scalability[58], and their potentially good performance with imbalance data classification[59]. We performed the training and optimisation (tuning) of these algorithms in each perspective using the *caret R Package*[60,61].

Classifiers training, optimisation and validation

**Class rebalancing techniques**: Our bipartite virus-mammal network is sparsely connected – roughly 0.23% of potential associations are documented in our input associations set. On average each virus in our dataset affected 3.45 mammals (~0.24%), and each mammalian host was affected by 4.41 viruses (~0.24%). This presented an imbalance in our data, whereby a small percentage of instances are positive (i.e. present a documented association between a virus and a mammal species). To attend to this issue, we deployed two class rebalancing techniques:

1. **SMOTE:** We deployed SMOTE - Synthetic Minority Over-sampling Technique[67,68] to rebalance the classes of our viral and mammalian models. SMOTE synthesises new minority class instances between existing minority instances using a flavour of k-nearest neighbour algorithm. The SMOTE algorithm then over-samples from the minority instances and under-samples from the majority class to create a balanced training set. SMOTE was applied to balance the training set of each mammalian and viral species model. It was not applied when drawing models' predictions or when assessing performance metrics as discussed below
2. **Random balanced under-sampling:** We trained our network perspective models (N=8×100) with balanced samples drawn at random (without replacement) from the set of all potential virus-mammal associations. Each sample comprised 2000 instances (1,000 positive (known) and 1,000 negative (true negative or unknown) virus-mammal associations.

**10-fold cross validation:** We performed 10-fold cross validation of our potential and constituent models (with various number of repeats – illustrated below). This validation methods works by splitting training data into 10 random samples, each sample is in turn is held out, and the model is trained on the remainder groups. The model's prediction for the existence or absence of the mammal-virus associations in the held-out group are used to construct confusion matrices and calculate an optimisation metric (here AUC). The optimisation metric is used to select best model in the validation process.

We repeated this cross validation for each of our constituent models as follows:

1. *Models trained in the viral and mammalian perspectives*:
   a. *Parameter tuning and classifier selection*: 10 repeats of 10-fold cross validation per model
   b. *Predictions and reported analyses*: 50 repeats of 10-fold cross validation per model. Results and metrics were obtained per repeat, and median, 0.05 and 0.95 quantiles for all scores were reported for predictions and metrics
2. *Models trained in the network perspective*: the 10-fold cross validation was repeated 100 times with balanced random samples drawn from the full network. Performance metrics, classifier selection, and predictions were obtained by testing each repeat against the full network. Median, 0.05 and 0.95 quantiles for all scores were reported.

| Model | Caret method | Base family | Summary |
|---|---|---|---|
| **Model Averaged Neural Network (avNNet)** | avNNet | Neural networks | The same neural network model is fit using different random number seeds. All resulting models are then used for prediction. To generate overall class probability, the probability outputs from all networks are averaged. |
| **Stochastic Gradient Boosting (GBM)** | gbm | Decision trees | Stochastic Gradient Boosting – GBM[57,62–64] fits a series of trees (weak classifiers) to random partition of the data, and aggregates the results sequentially (i.e. boosting). |
| **Random Forest (RF)** | ranger | | Random forests algorithm[65] construct an ensemble (termed forest) of decision trees grown randomly from the input data. The randomness is twofold: first, each tree is built from a random sample of the data. Second, at each tree node, a subset of features (input variables) is randomly selected to generate the best split. |
| **eXtreme Gradient Boosting (XGBoost)** | xgbTree | | Similarly to GBM, eXtreme Gradient Boosting (XGBoost)[66] constructs an ensemble (strong learner) from weak learners, typically decision trees. XGBoost has the additional advantages of speed; ease of use and parallelisation; and in many cases high predictive accuracy. |
| **Support Vector Machines with Radial basis kernel and Class Weights (SVM-RW)** | svmRadialWeights | Support vector machines | Support vector machines aim to find a hyperplane that best separates the features (predictors) into different domains (classes). Radial SVMs utilises Radial Basis Function Kernel (RBF), whereas polynomial SVMs adopts polynomial kernels. Linear SVMs uses linear kernels. SVMs with Class Weights – enjoys additional class-imbalance robustness as it applies penalty to miss classification, with weights inversely proportional to class frequency. |
| **Linear Support Vector Machines with Class Weights (SVM-LW)** | svmLinearWeights | | |
| **Support Vector Machines with Polynomial Kernel (SVM-P)** | svmPoly | | |
| **Naive Bayes (NB)** | naive_bayes | Bayesian | Naïve Bayes classifiers are a family of probabilistic classifiers centred upon Bayes Theorem. They assume that input features (virusal, mammalian and network features in our case) are independent – they contribute independently to probability to the outcome class (a virus-mammal association in our case), regardless of any correlation between the features. Features, however, are not always independent which is why the Naïve Bayes classifiers are labelled as naïve. |

**Table S4 - List of machine learning algorithms (supervised classifiers) used in our models.**

**Optimisation**: The hyper parameters of our viral and mammalian perspective models were tuned (optimised) via 10-fold cross validation (with 10 repeats) per each virus (or mammal) models. Hyper-parameters of best performing models were carried across to our bagging pipelines (50 repeats) which were used to produce median predictions (of class probabilities) and generate empirical confidence intervals. Models trained with motif-features (network-perspective) were optimised via 10-fold cross validation of each generated subsample (N=100).

We performed the training and tuning of our selected classifiers in each perspective using the *caret R Package*[60,61]. We adopted an adaptive resample approach[69] to tune the hyper-parameters of our models. This approach allows *caret* to adaptively resample the tuning parameter grid in a way that concentrates on values that are the in the neighbourhood of the optimal settings. Due to the large number of classifiers trained in our framework this adaptive approach allowed us to find optimal (or near optimal) values of the hyper-parameters of each included machine learning technique (table S4) without replaying on the nominal resampling process whereby all the tuning parameter combinations are computed for all the resamples before a choice is made about which parameters are good and which are poor.

#### Performance metrics

Confusion matrix

| | Detected | |
|---|---|---|
| Predicted | 1 | 0 |
| 1 | A | B |
| 0 | C | D |

| Measure | Formula | Meaning |
|---|---|---|
| Sensitivity (recall) | $\frac{A}{A+C}$ | Sensitivity is the percentage of actual positives (observed associations) that were correctly predicted. It indicates the percentage of 1s that was covered by the model. |
| Specificity | $\frac{D}{B+D}$ | Specificity is the percentage of negatives (here unknown associations, not necessarily true negative) that were correctly predicted |
| Precision | $\frac{A}{A+B}$ | Percentage of accurate predictions of the model |
| AUC | Area Under the ROC Curve | AUC is a threshold-independent measure of model predictive performance that is commonly used as a validation metric for host-pathogen predictive models[70,71]. |
| F1-score | $2 \times \frac{\text{Precision} \times \text{Recall}}{\text{Precision} + \text{Recall}}$ | Captures the harmonic mean of the precision and recall. it is often used with uneven class distribution. Our approach is relaxed with respect to false positives (unknown associations), hence the low F1-score recorded overall. However, in our selection process, where two classifiers produced similar AUC and TSS statistics, the best performing on F1-score was selected (conservative approach). |
| TSS | $\text{sensitivity} + \text{specificity} - 1$ | Use of AUC has been criticised for its insensitivity to absolute predicted probability and its inclusion of a priori untenable prediction [57,72], we also calculated the True Skill Statistic (TSS)[73]. |

**Table S5 - Measures utilised to assess the performance of our 10-fold cross-validated classifiers.** Entries highlighted in grey were not used in classifier selection.

#### Classifiers selection per perspective

We implemented two strategies to select constituent models of our three perspective, post training and optimising as per the previous subsection:

1. **Classifiers trained in the mammalian and viral feature spaces**: for each virus (or mammal) we selected the classifier which achieved the best performance metrics (table S5) overall (as derived from predictions). We were able to achieve this dynamic selection as we trained each classifier for each virus (or mammal) independently of other viruses (or mammals). We selected best performing classifiers based on their F1-Score, AUC, and True skill statistics – TSS (as defined in table S5 above).

2. **Classifiers trained in the motifs feature (network) space**: we selected the classifier which obtained the best test performance metric (median of 100 classifiers in each pipeline, applied to all 2,722,656 possible associations).

Held-out test set validation and performance metrics estimation

We estimated the performance of our framework and its constituent models by training these models on a stratified random sample comprising 85% of our input data (N=2,315,391 with 5,377 known virus-mammal associations), and then calculating performance metrics (table S5) on the held-out test set (N=407,265 known virus-mammal associations). Figures S4-S6 illustrates the validation process of each of our perspectives.

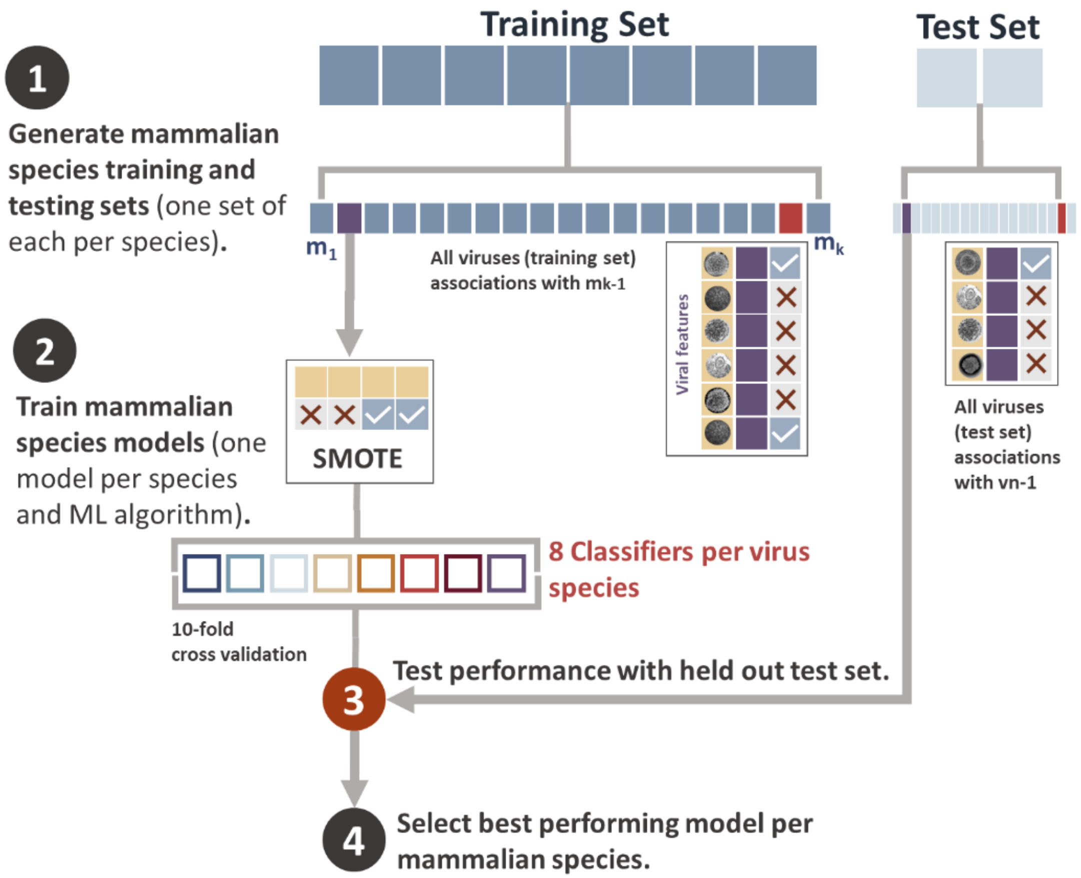


**Figure S4 – Training and validation of mammalian perspective models.** Training and test sets (85%, 15%) are split into (N=699) training and test sets, where each training/test pair contains associations of one mammalian species (e.g. humans) with all viruses in our input set (N=1,896). For each mammalian species, the training set is first balanced using SMOTE, then 8 classifiers are trained with this balanced set. These classifiers are then tested against the (unbalanced) test set, and the best performing (per each species) is carried forward to our multi-perspective framework.

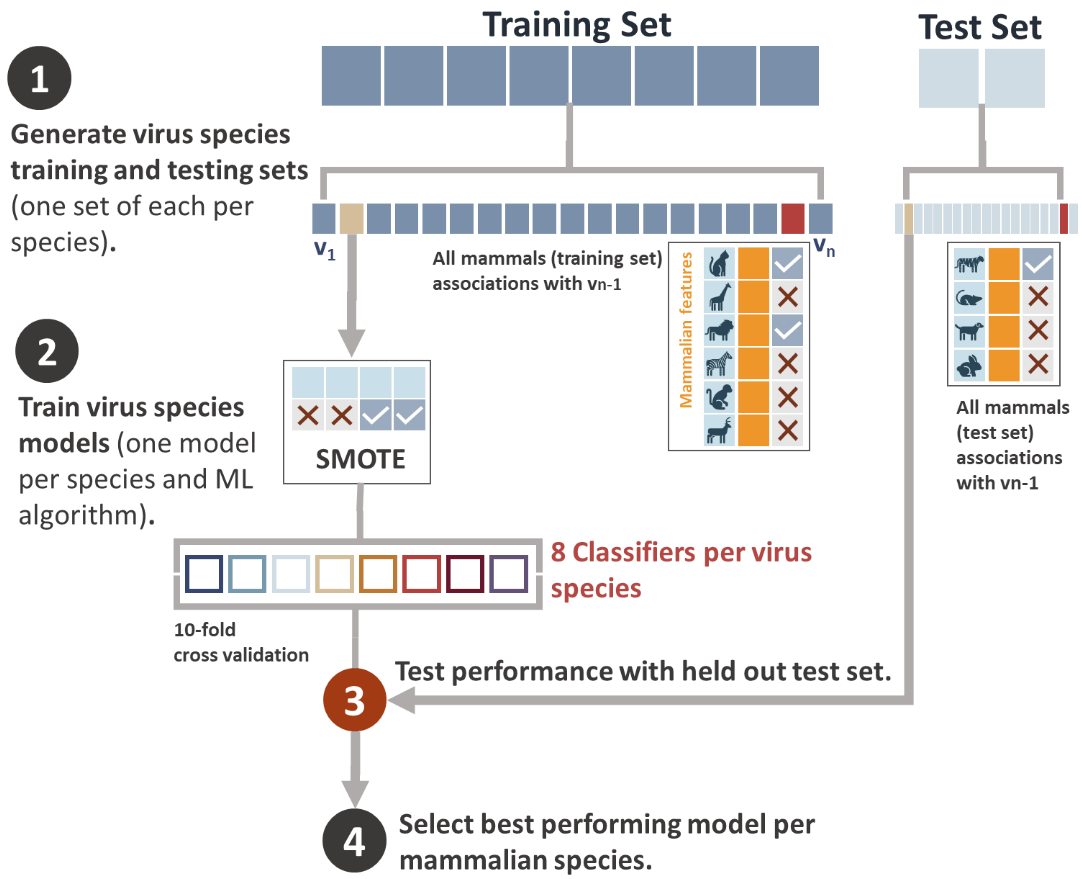


**Figure S5 – Training and validation of viral perspective models.** Training and test sets (85%, 15%) are split into (N=556) training and test sets, where each training/test pair contains associations of one mammalian species (e.g. humans) with all viruses in our input set (N=1,436). For each mammalian species, the training set is first balanced using SMOTE, then 8 classifiers are trained with this balanced set. These classifiers are then tested against the (unbalanced) test set, and the best performing (per each species) is carried forward to our multi-perspective framework.

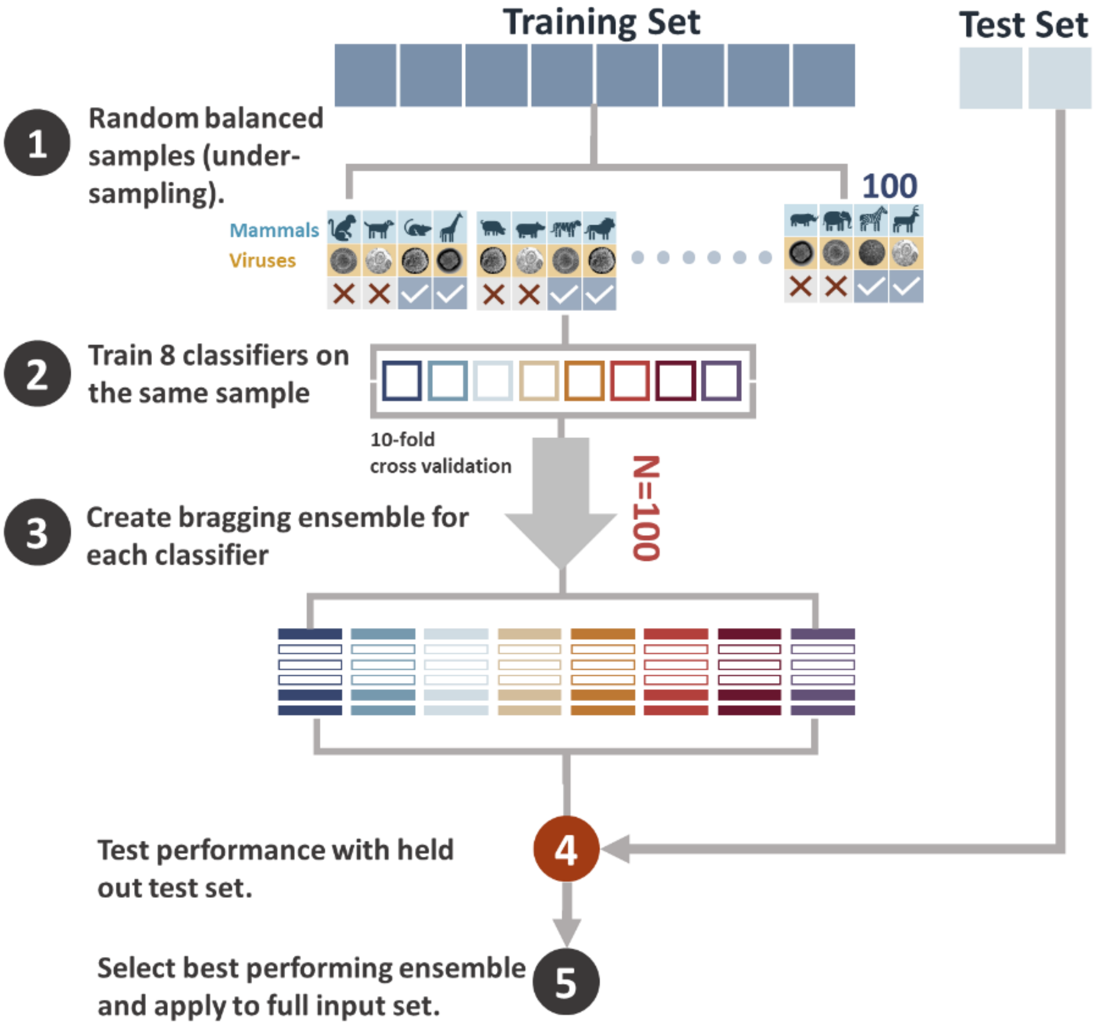


**Figure S6 – Training and validation of network perspective models.** 100 balanced samples (under-sampling) are drawn randomly from the training set (85% of all potential associations). Each sample (2,000 instances: 1,000 negative and 1,000 positive associations) is used to train 8 different classifiers (one of each of included algorithms - table S4). A Bragging (median prediction, probability) ensemble is then created for each algorithm. These ensembles are tested against the held-out test set (15%) and the best ensemble is carried forward to our multi-perspective framework.

#### Supplementary Results 1 – Systematic prediction of removed virus-mammal associations

In addition to validating each constituent model as described in the supplementary text and reported in manuscript and subsequent sections, we performed a systematic test to assess the ability of our framework to predict removed virus-mammal associations. This test differs from estimating performance metrics using a held-out test set (as per the procedures described in the supplementary text) in one important way: whereas the latter removes and tests the framework and its constituent model against larger amounts of held-out data (15% of all potential edges), the systematic test we describe below assesses our framework's ability to test potentially unseen virus-mammal associations by removing known (recorded in EID2) associations from our input set and recalculating all input parameters accordingly.

The test was performed systematically by removing one documented association (known to exist) between one virus and one mammal. The following input parameters were recalculated:

1. Recounting of all potential motifs as per the methods described above and in the main manuscript.
2. Recalculating the following mammalian features:
    a. Mean phylogenetic distance between focal mammal and known hosts of virus for each virus-mammal association.
    b. Mean ecological distance between focal mammal and known hosts of virus for each virus-mammal association.
3. Recalculating the following viral features:
    a. Mean phylogenetic distance between focal virus and viruses known to infect the associated mammal for each potential/known virus-mammal association
    b. Mean phylogenetic distance between focal virus and viruses known to infect the associated mammal
    c. Maximum phylogenetic distance between known hosts of virus.

Following these adjustments, we retrained all constituent models (mammalian, viral, and network perspectives) using 10-fold cross validation then attempted to predict the removed link.

We factored in differences in host-ranges of viruses, and variations in number of viruses detected per mammalian species by adopting the following processes:

1. **Viruses**: we ordered virus species by the number of unique mammalian hosts detected (in EID2), and second to fourth letter of virus name. We selected the first virus in each count category and ordered their hosts by the second to fourth letter of host species name (i.e. excluding genus). We selected the first host (in order) and removed the resulting host-virus association. Table S6 lists all removed associations from viruses' point of view.
2. **Mammals**: we ordered mammalian species by the number of unique virus species detected (in EID2), and second to fourth letter of species name (excluding genus). We selected the first mammalian species in each count category and ordered their viruses by the second to fourth letter of species name. We selected the first virus (in order) and removed the resulting host-virus association. Table S7 lists all removed associations from mammals' point of view.

**Table S6 - Systematic prediction of removed virus-mammal associations (viruses): virus-mammal associations removed from viruses' point of view**. No. is number of associations used to link with figure S7 (panel A – viruses), n = number of hosts per selected virus. Vote indicates if our framework predicted the removed association successfully (1) or not (0).

| No. | N | mammal | virus | vote | No. | N | mammal | virus | vote |
|---|---|---|---|---|---|---|---|---|---|
| 1 | 410 | *Homo sapiens* | tai forest ebolavirus | 1 | 23 | 23 | *Myodes glareolus* | dobrava-belgrade orthohantavirus | 1 |
| 2 | 134 | *Sus scrofa* | akabane orthobunyavirus | 1 | 24 | 22 | *Loxodonta africana* | rabies lyssavirus | 1 |
| 3 | 128 | *Bos taurus* | new jersey vesiculovirus | 1 | 25 | 21 | *Rousettus aegyptiacus* | rabies lyssavirus | 1 |
| 4 | 86 | *Pan troglodytes* | macaca mulatta polyomavirus 1 | 1 | 26 | 20 | *Odocoileus hemionus* | orf virus | 1 |
| 5 | 73 | *Macaca mulatta* | zika virus | 1 | 27 | 19 | *Dama dama* | rabies lyssavirus | 0 |
| 6 | 71 | *Equus caballus* | new jersey vesiculovirus | 1 | 28 | 18 | *Saimiri sciureus* | macaca mulatta polyomavirus 1 | 1 |
| 7 | 69 | *Ovis aries* | akabane orthobunyavirus | 1 | 29 | 17 | *Papio hamadryas* | baboon orthoreovirus | 1 |
| 8 | 65 | *Canis lupus familiaris* | betacoronavirus 1 | 1 | 30 | 16 | *Eptesicus serotinus* | bat astrovirus | 1 |
| 9 | 54 | *Felis catus* | rabies lyssavirus | 1 | 31 | 15 | *Microtus agrestis* | rabies lyssavirus | 0 |
| 10 | 53 | *Rattus norvegicus* | rat astrovirus | 1 | 32 | 14 | *Syncerus caffer* | rinderpest morbillivirus | 1 |
| 11 | 47 | *Mus musculus* | thailand orthohantavirus | 1 | 33 | 13 | *Callithrix jacchus* | rabies lyssavirus | 0 |
| 12 | 45 | *Macaca fascicularis* | tai forest ebolavirus | 1 | 34 | 12 | *Chlorocebus sabaeus* | pegivirus a | 1 |
| 13 | 44 | *Eidolon helvum* | rabies lyssavirus | 1 | 35 | 11 | *Giraffa camelopardalis* | betacoronavirus 1 | 1 |
| 14 | 40 | *Miniopterus schreibersii* | bat astrovirus 1 | 1 | 36 | 10 | *Pteropus scapulatus* | rabies lyssavirus | 1 |
| 15 | 35 | *Chlorocebus aethiops* | measles morbillivirus | 1 | 37 | 9 | *Lontra canadensis* | rabies lyssavirus | 0 |
| 16 | 34 | *Zalophus californianus* | sea lion mastadenovirus a | 1 | 38 | 8 | *Camelus bactrianus* | rotavirus a | 1 |
| 17 | 33 | *Equus asinus* | betacoronavirus 1 | 1 | 39 | 7 | *Macaca radiata* | kyasanur forest disease virus | 1 |
| 18 | 32 | *Cervus elaphus* | rabies lyssavirus | 1 | 40 | 6 | *Vulpes lagopus* | rabies lyssavirus | 1 |
| 19 | 30 | *Rattus rattus* | rat astrovirus | 1 | 41 | 5 | *Tylonycteris pachypus* | bat circovirus | 1 |
| 20 | 28 | *Rhinolophus ferrumequinum* | bat bocavirus | 1 | 42 | 4 | *Saguinus labiatus* | hepatovirus a | 1 |
| 21 | 26 | *Rangifer tarandus* | orf virus | 1 | 43 | 3 | *Plecotus rafinesquii* | rabies lyssavirus | 1 |
| 22 | 24 | *Capreolus capreolus* | rotavirus a | 1 | | | | | |

**Table S7 - Systematic prediction of removed virus-mammal associations (mammals): virus-mammal associations removed from mammals' point of view**. No. is number of associations used to link with figure S7 (panel B – mammals), N = number of viruses per selected mammal. Vote indicates if our framework predicted the removed association successfully (1) or not (0).

| No. | N | mammal | virus | vote | No. | n | mammal | virus | vote |
|---|---|---|---|---|---|---|---|---|---|
| 1 | 682 | *Chrysocyon brachyurus* | rabies lyssavirus | 1 | 24 | 26 | *Rattus tanezumi* | rodent astrovirus | 1 |
| 2 | 97 | *Callithrix jacchus* | monkeypox virus | 1 | 25 | 25 | *Cervus elaphus* | respiratory syncytial virus | 1 |
| 3 | 78 | *Chrysocyon brachyurus* | canine morbillivirus | 1 | 26 | 24 | *Equus caballus* | saint louis encephalitis virus | 1 |
| 4 | 62 | *Vulpes macrotis* | carnivore protoparvovirus 1 | 1 | 27 | 23 | *Oligoryzomys flavescens* | andes orthohantavirus | 1 |
| 5 | 57 | *Lama glama* | rotavirus a | 1 | 28 | 22 | *Oligoryzomys flavescens* | andes orthohantavirus | 1 |
| 6 | 56 | *Equus caballus* | west nile virus | 1 | 29 | 21 | *Tayassu pecari* | suid alphaherpesvirus 1 | 0 |
| 7 | 55 | *Artibeus planirostris* | influenza a virus | 0 | 30 | 20 | *Boselaphus tragocamelus* | small ruminant morbillivirus | 1 |
| 8 | 50 | *Dama dama* | tick-borne encephalitis virus | 1 | 31 | 19 | *Acinonyx jubatus* | felid alphaherpesvirus 1 | 1 |
| 9 | 48 | *Lama glama* | pestivirus a | 1 | 32 | 18 | *Pteropus giganteus* | pegivirus b | 0 |
| 10 | 45 | *Lama pacos* | bluetongue virus | 1 | 33 | 17 | *Macaca cyclopis* | macaca mulatta polyomavirus 1 | 1 |
| 11 | 44 | *Pteropus scapulatus* | japanese encephalitis virus | 1 | 34 | 16 | *Equus caballus* | deltapapillomavirus 4 | 1 |
| 12 | 43 | *Hipposideros caffer* | bat paramyxovirus | 1 | 35 | 14 | *Ovis dalli* | orf virus | 1 |
| 13 | 42 | *Equus caballus* | ovine gammaherpesvirus 2 | 1 | 36 | 13 | *Chlorocebus sabaeus* | pegivirus a | 1 |
| 14 | 41 | *Pteropus scapulatus* | dengue virus | 1 | 37 | 12 | *Homo sapiens* | dobrava-belgrade orthohantavirus | 1 |
| 15 | 40 | *Equus caballus* | eastern equine encephalitis virus | 1 | 38 | 11 | *Equus caballus* | akabane orthobunyavirus | 1 |
| 16 | 39 | *Cynopterus brachyotis* | bat coronavirus | 1 | 39 | 10 | *Vulpes vulpes* | rabbit hemorrhagic disease virus | 0 |
| 17 | 38 | *Equus caballus* | orthohepevirus a | 1 | 40 | 9 | *Akodon azarae* | argentinian mammarenavirus | 1 |
| 18 | 36 | *Boselaphus tragocamelus* | alcelaphine gammaherpesvirus 1 | 0 | 41 | 8 | *Macaca radiata* | kyasanur forest disease virus | 1 |
| 19 | 35 | *Saimiri sciureus* | yellow fever virus | 1 | 42 | 7 | *Mandrillus sphinx* | macacine gammaherpesvirus 5 | 1 |
| 20 | 34 | *Cercopithecus nictitans* | lymphocryptovirus 1 | 1 | 43 | 6 | *Homo sapiens* | bhanja virus | 1 |
| 21 | 32 | *Hipposideros abae* | rift valley fever phlebovirus | 1 | 44 | 5 | *Zalophus californianus* | seal parapoxvirus | 1 |
| 22 | 31 | *Puma yagouaroundi* | alphacoronavirus 1 | 1 | 45 | 4 | *Rattus rattus* | thailand orthohantavirus | 1 |
| 23 | 30 | *Equus caballus* | cowpox virus | 1 | 46 | 3 | *Peromyscus leucopus* | rabies lyssavirus | 1 |

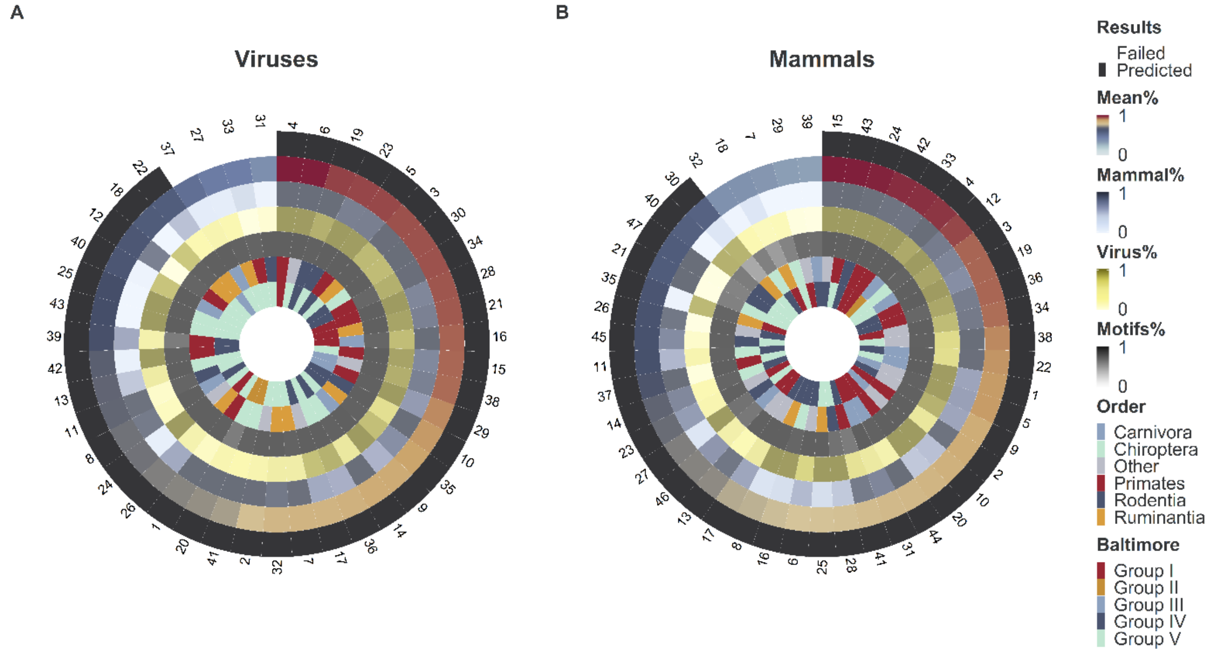


**Figure S7 – Results of systematic prediction of removed virus-mammal associations** . Circles represent the following in order: whether the leave out test succeeded in predicting the removed interaction (black) or not (white); the mean probability across the three perspectives; probabilities derived from the mammalian perspective; probabilities derived from the viral perspective; probabilities derived from the network (motifs) perspective; the order of the host; and Baltimore classification of the virus**. Panel A – results of viruses (interactions removed for each virus selected in table S6. Panel B – results of mammals (interactions removed for each virus selected in table S7**.

**Supplementary Results 2 – The mammalian perspective**

#### Classifier selection

We investigated the effect viral traits has on the distribution of 1,896 virus species in each of the 699 mammalian species which we found to host at least 2 virus species using 8 supervised classification algorithms (table S4). We optimised (tuned) the hyper-parameters of each classifier using 10-fold cross validation repeated 10 times, with SMOTE class rebalancing technique. Classifier selection per mammal was achieved using the strategy outlined in Supplementary Note 4. Selected classifiers (i.e. algorithms) and their hyper-parameters (best tunes) were carried across to bagging pipelines, whereby each classifier was retrained with 10-fold cross validation repeated 50 times (with same tuning parameters), to attend to uncertainties in our sampling regiments and the cross-validation processes, and to generate empirical confidence intervals. Predictions were generated using median probability of selected classifiers pipelines, 90% CI were also obtained and were integrated into our framework pipeline.

**Validation**: When taking median predictions of our 10-fold cross validated mammalian perspective models (trained with all available data for each mammal, via cross validation), our mammalian perspective pipelines achieved an AUC = 0.916, F1-score = 0.201, and TSS = 0.832. When trained with 85% of available data our mammalian perspective performance was as follows: AUC=0.816 F1-score = 0.104 and TSS=0.632. Our mammalian perspective performance on the held-out test set (15%) was as follows: AUC=0.807 F1-score = 0.089 and TSS=0.614.

#### New mammal-virus associations obtained from our mammalian perspective models

When using viral traits to predict associations between mammals and their viruses the results of our selected models (previous section) suggested that 41,537 (median, 90% CI [4275, 238971]) unknown associations could be missing from our original dataset. Our mammalian perspective models suggested 33.24 [6.93, 170.82] viruses per host (~8.25-fold increase [~1.36, ~37.6]), with median 28.93 [2.98, 166.41] new viruses per host. Figure S10 illustrates the results obtained from our mammalian perspectives.

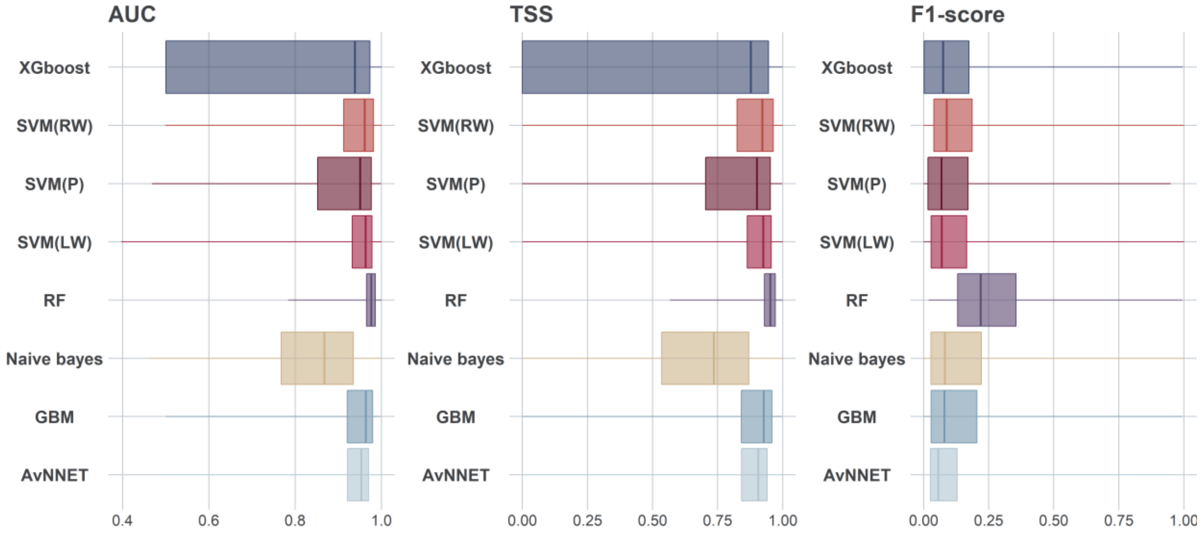


**Figure S8 - Comparison of machine learning algorithms trained in viral feature space (mammalian perspective).** Boxplots show the median, quartiles and range of performance metrics for 8 algorithms from 10-fold cross validation, each trained with same training sets per each mammal (N=699).

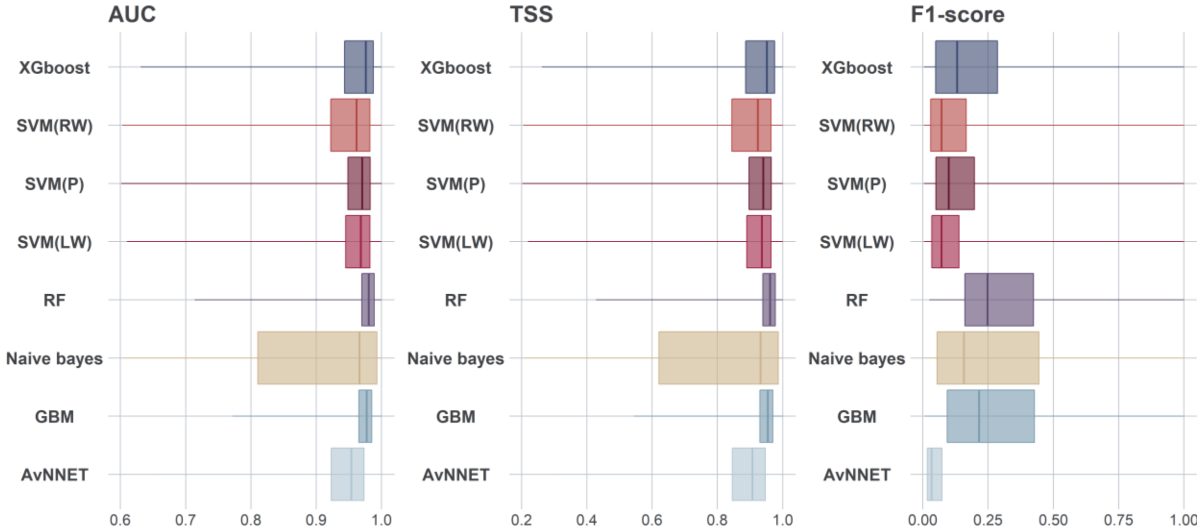


**Figure S9 - Selected classifiers trained in viral feature space (mammalian perspective).** Boxplots show the median, quartiles, and range of performance metrics for 8 algorithms from 50 runs with 10-fold cross validation of best performing tuned classifiers (Figure S8) per each mammal (n=699). Percentage of selected classifiers (number of times a classification algorithm was selected as best performing for a given mammalian species divided by number of times the algorithm was run in total) was as follows: avNNet=08.96%, GBM =10.90%, RF=11.64%, XGBoost=09.25%, SVM-RW=23.43%, SVM-LW=15.52%, SVM-P=11.19%, NB=09.10%.

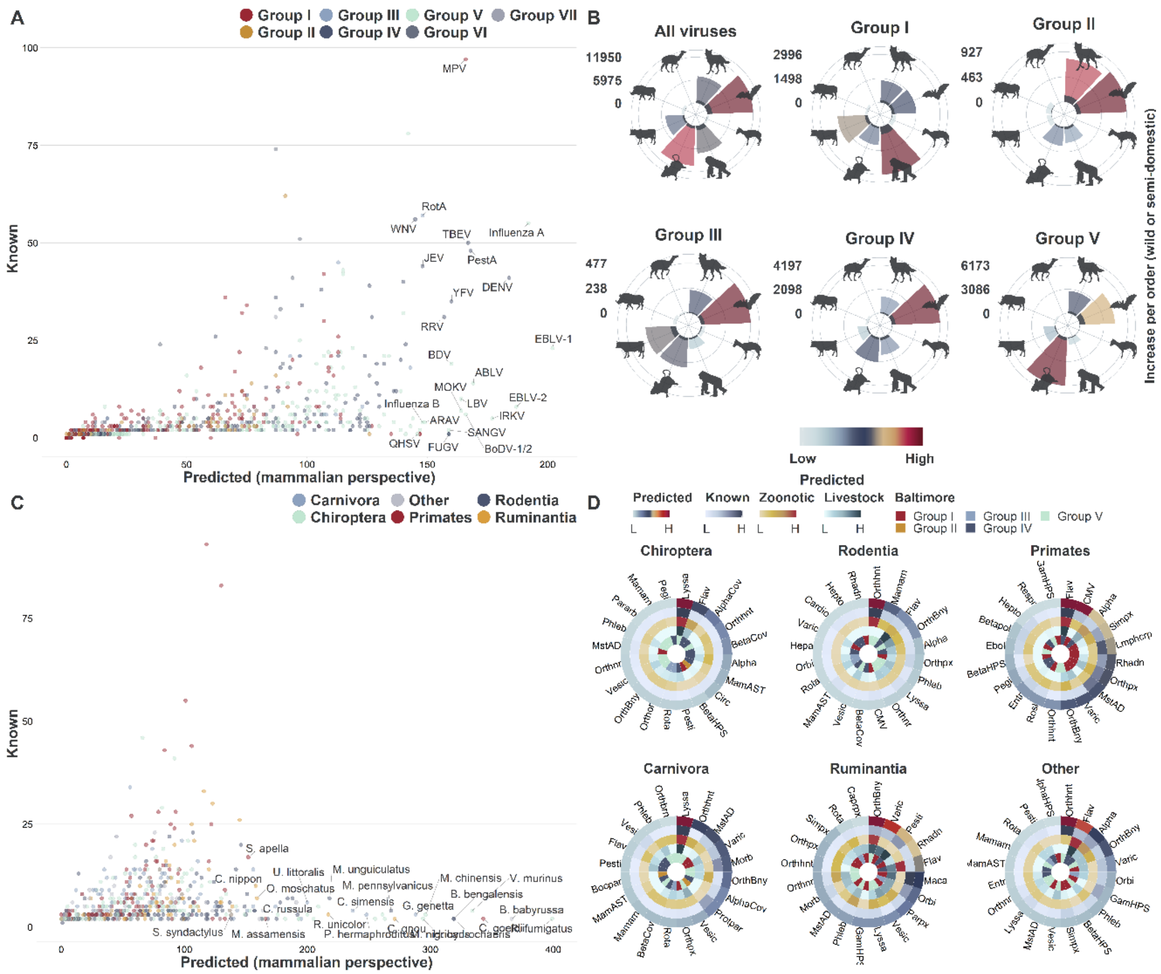


**Figure S10 - results of our mammalian perspective models. Panel A – changes in mammalian host range of viruses from known (in EID2) to predicted (by mammalian perspective models – top 30 are labelled)**. Rabies lyssavirus was excluded from panel A to allow for better visualisation. **Panel B – predicted host-virus associations per order between viruses and wild and semi-domesticated mammals**. We selected the following orders to represent (clockwise): Carnivora, Chiroptera, Perissodactyla, Primates, Rodentia, Ruminantia, Suina and Tylopoda. Dark grey bars indicate detected interaction (in EID2). **Panel C – changes in number of viruses per wild or semi-domesticated mammalian host species from detected (in EID2) to predicted** (by mammalian perspective models – top 30 are labelled). **Panel D – Top 15 genera (by number of predicted wild or semi-domesticated mammalian host species) in selected orders (Other indicated results for all orders not included in the first five circles)**. Each order figure comprises the following circles (from outside to inside): 1) number of hosts predicted to have an association with the viral genus; 2) number of hosts detected to have association; 3) number of hosts predicted to harbour viral zoonotic (i.e. predicted to share at least one virus species with humans); 4) number of hosts predicted to share viruses with domesticated livestock (domesticated mammals in orders Perissodactyla, Ruminantia, Suina and Tylopoda); and 5) Baltimore classification of the selected genera.

#### Supplementary Results 3 – the viral perspective

Classifier selection

We examined the effect of mammalian traits on the distribution of each of the 556 virus species for which we found more than one mammalian host in 1,436 terrestrial mammals using the same classifier selection strategy as the previous section. Predictions were generated using median probability of selected classifiers pipelines, 90% CI were also obtained and were integrated into our pipeline.

**Validation**: When taking median predictions of our 10-fold cross validated viral perspective models (trained with all available data for each mammal, via cross validation), our mammalian perspective pipelines achieved an AUC = 0.867, F1-score = 0.289, and TSS = 0.734. When trained with 85% of available data our viral perspective performance was as follows: AUC=0.800 F1-score = 0.150 and TSS=0.600. Our viral perspective performance on the held-out test set (15%) was as follows: AUC=0.797 F1-score = 0.150 and TSS=0.594.

New links (mammalian host-virus associations)

When using mammalian features to predict potential associations with multi-host viruses, the results of our selected models indicated that 21,352 (median, 90% CI [2536, 95630]) unknown associations could be missing from the original bipartite network. Our mammalian perspective models suggested a median 14.56 [4.48, 53.78] hosts per virus (~2.96 [~1.15, ~11.84]), with median 11.26 [1.34, 50.44] new hosts per virus. Figure S13 illustrates the results obtained from our viral perspective models.

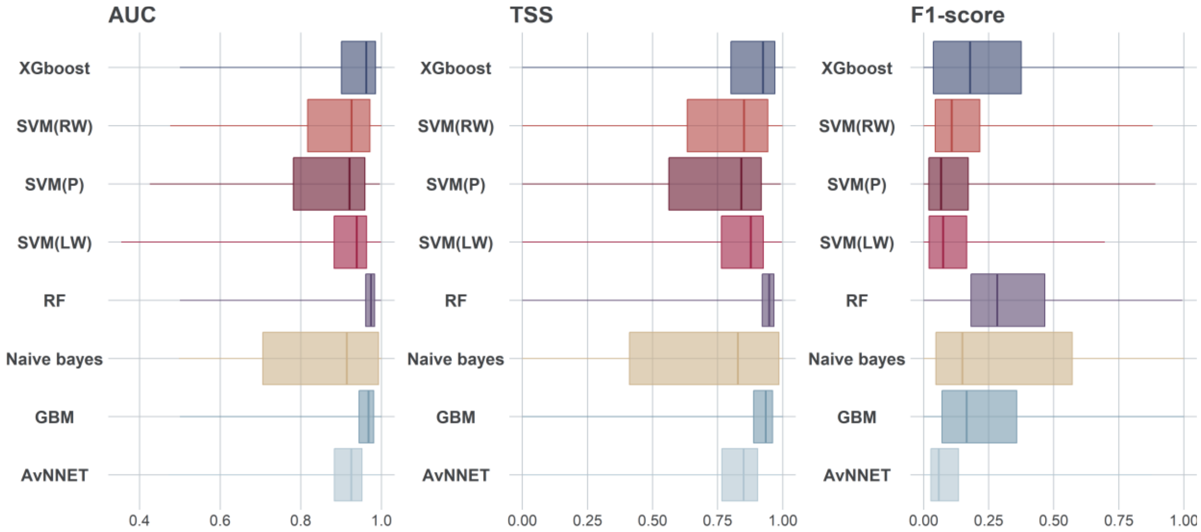


**Figure S11 - Comparison of machine learning algorithms trained in mammalian feature space (viral perspective)**. Boxplots show the median, quartiles, and range of performance metrics for 8 algorithms from 10-fold cross validation, each trained with same training sets per each virus (n=556).

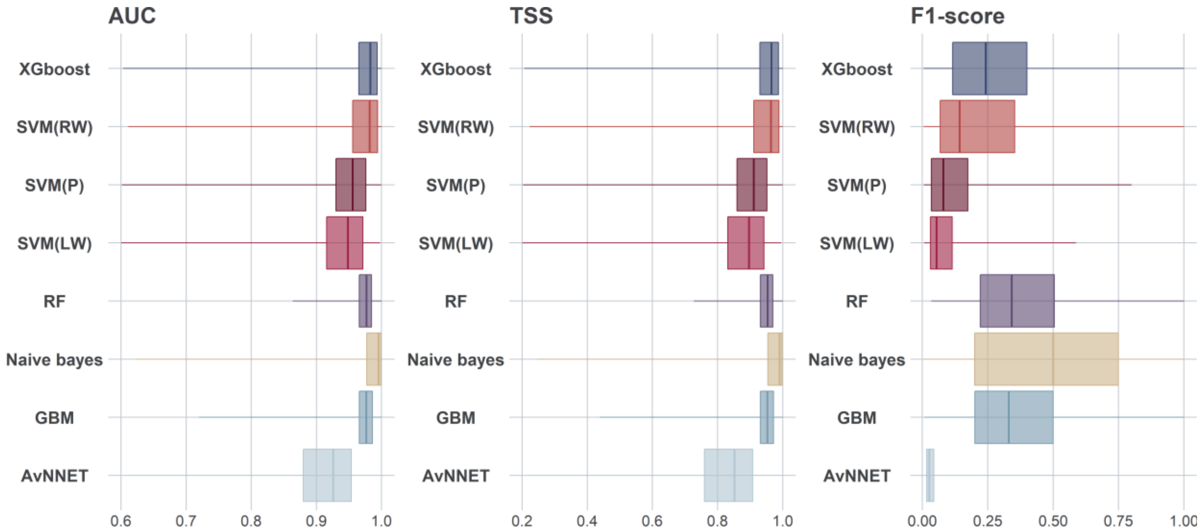


**Figure S12 - Selected classifiers trained in mammalian feature space (viral perspective).** Boxplots show the median, quartiles, and range of performance metrics for 8 algorithms from 50 runs with 10-fold cross validation of best performing tuned classifiers (Figure S11) per each virus (n=556). Percentage of selected classifiers (number of times a classification algorithm was selected as best performing for a given virus species divided by number of times the algorithm was run in total) was as follows: avNNet =6.18%, GBM =16.6%, RF=13.13%, XGBoost=20.46%, SVM-RW=14.48%, SVM-LW=4.63%, SVM-P=5.02%, NB=19.5%.

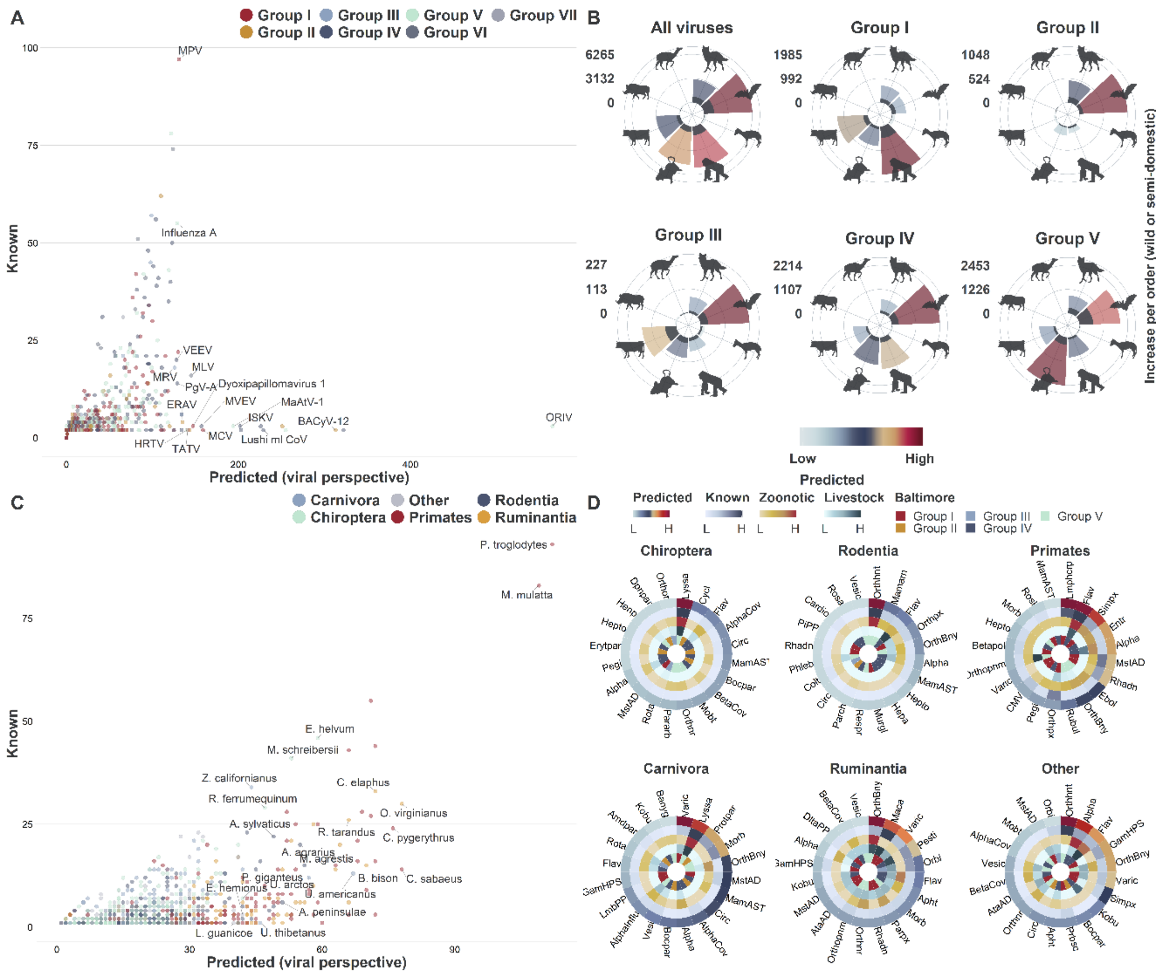


**Figure S13 - results of our viral perspective models. Panel A – changes in mammalian host range of viruses from known (in EID2) to predicted (by viral perspective models – top 30 are labelled).** Rabies lyssavirus was excluded from panel A to allow for better visualisation. **Panel B – predicted host-virus associations per order between viruses and wild and semi-domesticated mammals**. We selected the following orders to represent (clockwise): Carnivora, Chiroptera, Perissodactyla, Primates, Rodentia, Ruminantia, Suina and Tylopoda. Dark grey bars indicate detected interaction (in EID2). **Panel C – changes in number of viruses per wild or semi-domesticated mammalian host species from detected (in EID2) to predicted** (by viral perspective models – top 30 are labelled). **Panel D – Top 15 genera (by number of predicted wild or semi-domesticated mammalian host species) in selected orders (Other indicated results for all orders not included in the first five circles)**. Each order figure comprises the following circles (from outside to inside): 1) number of hosts predicted to have an association with the viral genus; 2) number of hosts detected to have association; 3) number of hosts predicted to harbour viral zoonotic (i.e. predicted to share at least one virus species with humans); 4) number of hosts predicted to share viruses with domesticated livestock (domesticated mammals in orders Perissodactyla, Ruminantia, Suina and Tylopoda); and 5) Baltimore classification of the selected genera.

#### Supplementary Results 4 – The network perspective

Classifier selection

We performed full network prediction (all possible edges, N = 2,722,656) using our trained classifiers (as per the sampling and aggregating strategy listed in main manuscript and Supplementary Note 4). We assessed the results of these classifiers (including unseen data) using the metrics outlined in Supplementary Note 4. Following this, SVM(RW) (Support Vector Machine with RBF kernel and class weights) emerged as the best classifier overall.

**Validation**: When taking median predictions of our 10-fold cross validated network perspective models (trained with all available virus-mammal associations, via cross validation), our network perspective pipelines achieved an AUC = 0.959, F1-score = 0.139, and TSS = 0.918. When trained with 85% of available data our viral perspective performance was as follows: AUC=0.927 F1-score = 0.300 and TSS=0.856. Our network perspective performance on the held-out test set (15%) was as follows: AUC=0.925 F1-score = 0.138 and TSS=0.850.

New links (mammalian host-virus associations)

When inferring association between mammals and their viruses using motifs as features, our models suggested 76,081 (median, 90% CI [27738, 205814]) unknonw associations could be missing from the original bipartite network. Our network perspective models indicated a median 57.15 [23.22, 147.62] viruses per mammalian host (median fold increase ~14.12 [~4.64, ~34.29]), with 52.98 [19.32, 143.32] new viruses per host. Conversely, they suggested a median 43.29 [17.59, 111.81] mammalian hosts per virus (~7.58 [~2.75, ~25.25]), with 40.13 [14.63, 108.55] new hosts per virus. Figure S15 illustrates the results obtained from our networks perspective models.

Variable importance – Motifs selection

As stated in the main manuscript, we did not filter motifs prior to feeding them to the machine learning algorithms. Figure S16 illustrates the relative influence of motifs count per edge (simply motifs) in our selected models – SVM (RW).

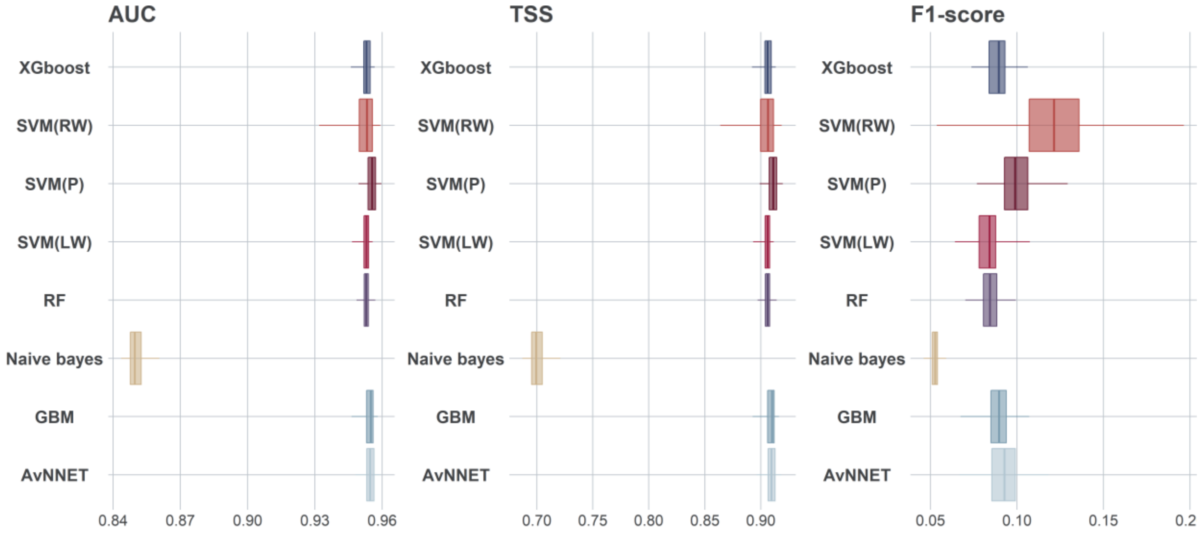


**Figure S14 - Comparison of machine learning algorithms trained in mammalian feature space (viral perspective).** Boxplots show the median, quantiles, and range of performance metrics for 8 algorithms from 10-fold cross validation (100 repeats), results derived from applying each trained model to the set of possible mammal virus interactions (N=2,722,656).

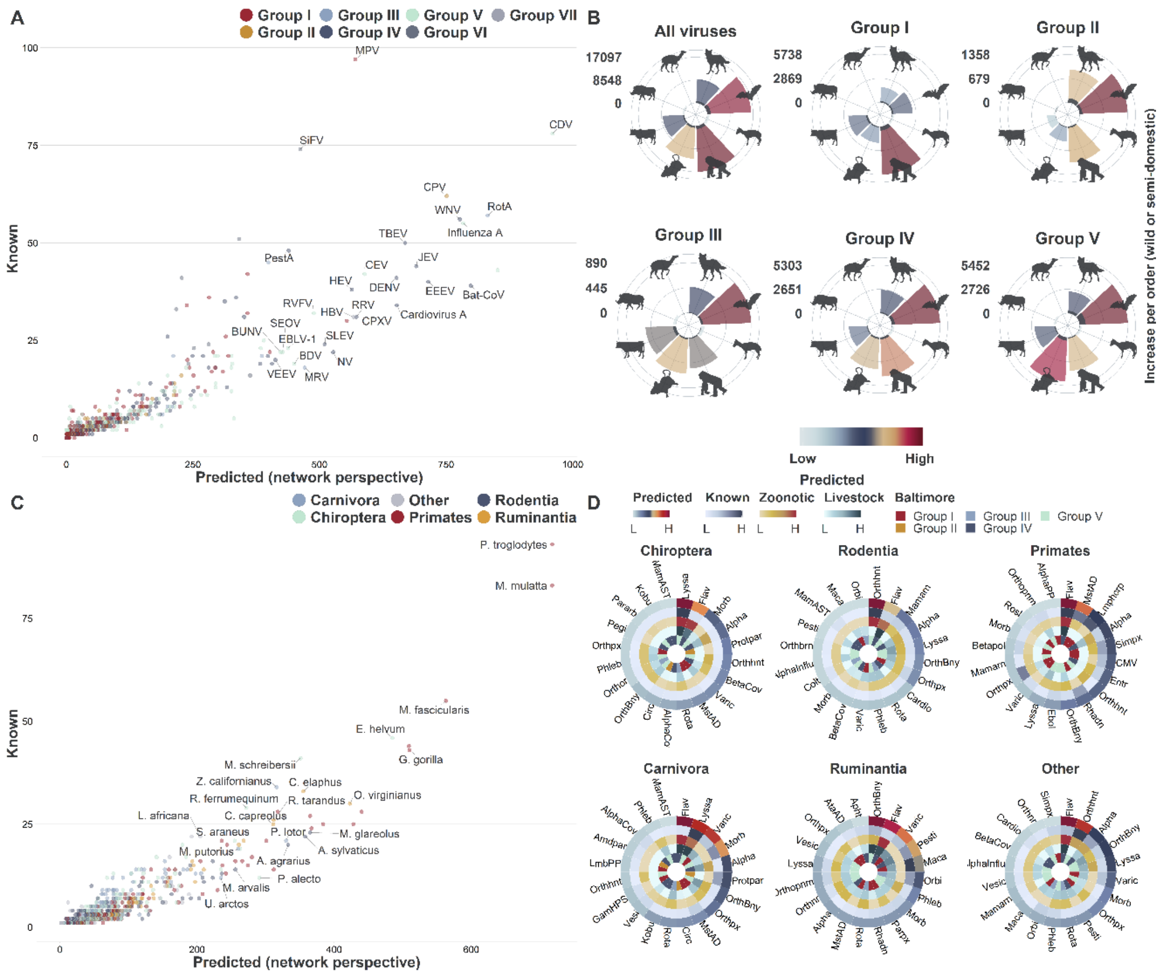


**Figure S15 - results of our network perspective models. Panel A – changes in mammalian host range of viruses from known (in EID2) to predicted (by network perspective models – top 30 are labelled).** Rabies lyssavirus was excluded from panel A to allow for better visualisation. **Panel B – predicted host-virus associations per order between viruses and wild and semi-domesticated mammals.** We selected the following orders to represent (clockwise): Carnivora, Chiroptera, Perissodactyla, Primates, Rodentia, Ruminantia, Suina and Tylopoda. Dark grey bars indicate detected interaction (in EID2). **Panel C – changes in number of viruses per wild or semi-domesticated mammalian host species from detected (in EID2) to predicted** (by network perspective models – top 30 are labelled). **Panel D – Top 15 genera (by number of predicted wild or semi-domesticated mammalian host species) in selected orders (Other indicated results for all orders not included in the first five circles).** Each order figure comprises the following circles (from outside to inside): 1) number of hosts predicted to have an association with the viral genus; 2) number of hosts detected to have association; 3) number of hosts predicted to harbour viral zoonotic (i.e. predicted to share at least one virus species with humans); 4) number of hosts predicted to share viruses with domesticated livestock (domesticated mammals in orders Perissodactyla, Ruminantia, Suina and Tylopoda); and 5) Baltimore classification of the selected genera.

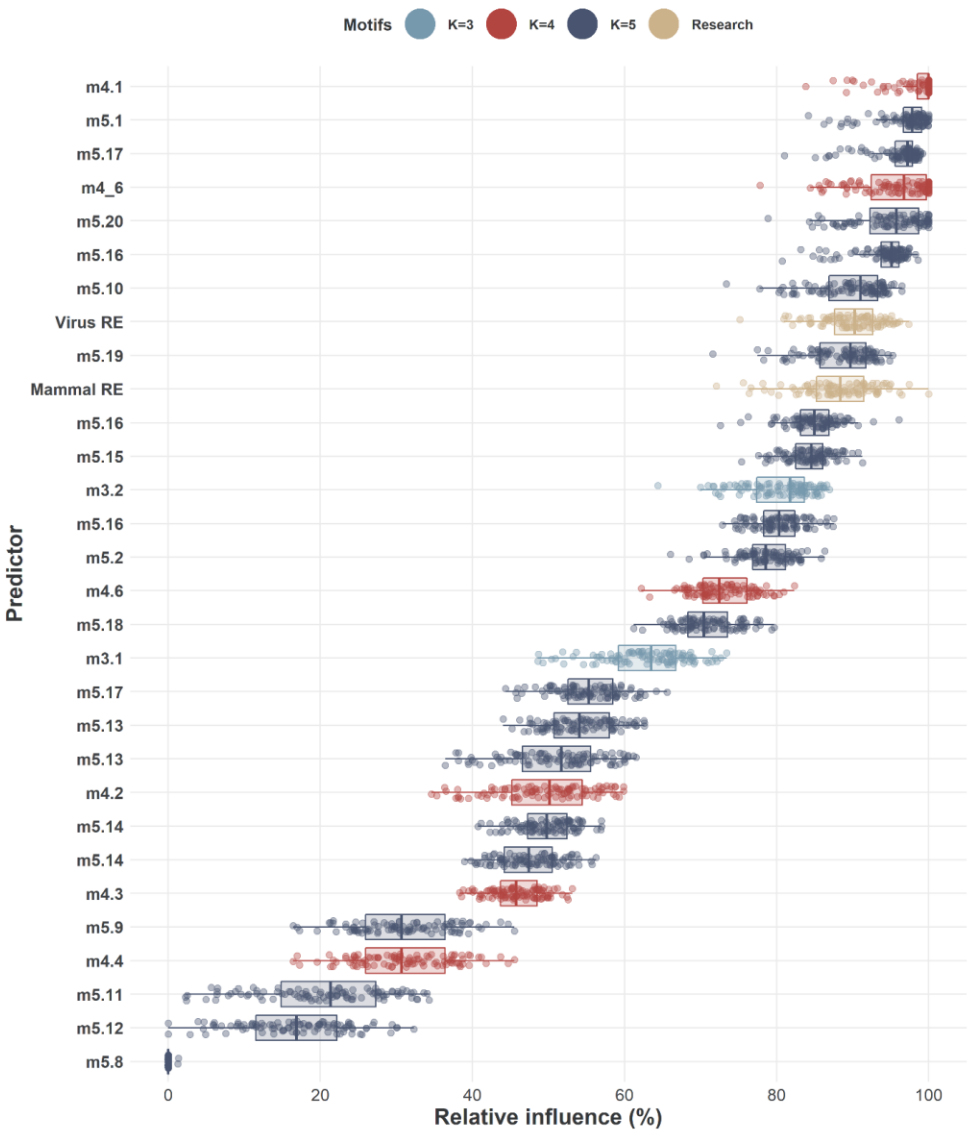


**Figure S16 - Relative contribution (importance) of motif-features (variables) to the AUC of our SVM-RW models trained with random balanced subsampling (n=100 times) in with our "potential" motif-features as predictors of unknown virus-host association (new links)**. Motifs (subgraphs) are coloured by the number of nodes (K=3, 4, 5) and correspond to figure 1 in the main manuscript. Research effort into both viruses and mammals was added as independent variables in our motifs models (coloured in yellow).

#### Supplementary Results 5 – Additional Results

Difference in performance metrics between training and testing sets

**Table S8– Differences in performance metrics between training and testing sets at full model and individual perspective levels.**

| Perspective | | AUC | F1-Score | TSS |
|---|---|---|---|---|
| Full model | Training | 0.972 | 0.322 | 0.944 |
| | Testing | 0.956 | 0.310 | 0.915 |
| Mammalian | Training | 0.916 | 0.138 | 0.832 |
| | Testing | 0.890 | 0.115 | 0.781 |
| Viral | Training | 0.867 | 0.194 | 0.735 |
| | Testing | 0.839 | 0.181 | 0.678 |
| Network | Training | 0.952 | 0.192 | 0.904 |
| | Testing | 0.947 | 0.190 | 0.893 |

#### Viruses found in human

Our multi-perspective framework generates predictions for each potential virus-mammal association (N=2,722,656 between 1,896 viruses and 1,436 terrestrial mammals). Here we highlight results for humans. In addition to 425 virus species known to affect humans, our model predicted 16 undocumented (in EID2) virus species that could potentially be found in humans. Table S9 presents these viruses.

**Table S9– Predicted viruses that could potentially be found in humans (not documented in input dataset)**. *Baltimore* is Baltimore classification (7 groups I to VII). *Mammalian* lists probabilities drawn from *median* mammalian perspective (Homo sapiens model trained with viral traits). *Viral* lists *median* probabilities drawn for humans from viral perspective (16 virus models trained with mammalian traits). *Network* lists probabilities drawn from network perspective (trained with motif features). *Mean:* is average probability across the three perspectives. Voting was drawn for each perspective (1 if probability 0.5).

| Virus species | Baltimore | Mammalian | Viral | Network | Mean |
|---|---|---|---|---|---|
| murid betaherpesvirus 1 | I | 0.003 | 0.541 | 0.999 | 0.514 |
| cercopithecine alphaherpesvirus 9 | I | 0.042 | 0.781 | 0.999 | 0.607 |
| lymphocryptovirus 1 | I | 0.073 | 1.000 | 0.999 | 0.691 |
| deltapapillomavirus 4 | I | 0.002 | 0.994 | 0.997 | 0.665 |
| bat rotavirus | III | 0.005 | 0.601 | 0.988 | 0.531 |
| bat hepatovirus | IV | 0.006 | 0.979 | 0.973 | 0.653 |
| primate astrovirus | IV | 0.045 | 0.618 | 0.999 | 0.554 |
| enterovirus j | IV | 0.158 | 0.707 | 0.999 | 0.621 |
| enterovirus f | IV | 0.007 | 0.708 | 0.996 | 0.570 |
| mamastrovirus 5 | IV | 0.574 | 0.224 | 0.998 | 0.599 |
| aravan lyssavirus | V | 0.004 | 0.519 | 0.955 | 0.493 |
| beilong jeilongvirus | V | 0.001 | 0.518 | 0.990 | 0.503 |
| coastal plains tibrovirus | V | 0.008 | 0.627 | 0.996 | 0.544 |
| influenza d virus | V | 0.019 | 0.902 | 0.998 | 0.640 |
| simian immunodeficiency virus | VI | 0.459 | 0.696 | 0.999 | 0.718 |
| simian retrovirus 5 | VI | 0.021 | 0.575 | 0.997 | 0.531 |

#### Rare viruses and rarely studied mammals

Our multi-perspective framework can capture and expand associations of both rare viruses, and understudied mammalian species, due to separation of perspectives. If a virus is rare, the framework would capture potential hosts via the network and mammalian perspectives. Similarly, if a mammalian species is rarely studied, then the framework would still capture viruses potentially found in this mammalian species via the network and viral perspectives. In relation to rare viruses and understudied mammalian species, our framework was able to expand the host range of rare viruses (n=1,450) from 1,619 to 4,174 (~ 2.16 average increase per rare virus). Virus range of rare mammals (n=954) was also increased from 1,150 to 4,318 (~3.21 average fold increase per host).
